# Defining the activation profile and fate trajectory of adult Scleraxis-lineage cells during tendon healing by combining lineage tracing and spatial transcriptomics

**DOI:** 10.1101/2021.06.02.446663

**Authors:** Jessica E. Ackerman, Katherine T. Best, Samantha N. Muscat, Chia-Lung Wu, Alayna E. Loiselle

**Affiliations:** Center for Musculoskeletal Research, Department of Orthopaedics & Rehabilitation, University of Rochester Medical Center, Rochester, NY

## Abstract

The tendon healing process is regulated by the coordinated interaction of multiple cell types and molecular processes. However, these processes are not well-defined leading to a paucity of therapeutic approaches to enhance tendon healing. Scleraxis-lineage (Scx^Lin^) cells are the major cellular component of adult tendon and make time-dependent contributions to the healing process. Prior work from our lab and others suggests heterogeneity within the broader Scx^Lin^ population over the course of tendon healing; therefore delineating the temporal and spatial contributions of these cells is critical to understanding and improving the healing process. In the present study we utilize lineage tracing of the adult aScx^Lin^ population to determine whether these cells undergo cellular activation and subsequent myofibroblast differentiation, which is associated with both proper healing and fibrotic progression in many tissues. We show that adult aScx^Lin^ cells undergo transient activation in the organized cellular bridge at the tendon repair site, contribute to the formation of an organized neotendon, and contribute to a persistent myofibroblast population in the native tendon stubs. The mechanisms dictating this highly specialized spatial response are unknown. We therefore utilized spatial transcriptomics to better define the spatio-molecular program of tendon healing. Integrated transcriptomic analyses across the healing time-course identifies five distinct molecular regions, including key interactions between the inflammatory bridging tissue and highly reactive tendon tissue at the repair site, with adult Scx^Lin^ cells being a central player in the transition from native tendon to reactive, remodeling tendon. Collectively, these data provide important insights into both the role of adult Scx^Lin^ cells during healing as well as the molecular mechanisms that underpin and coordinate the temporal and spatial healing phenotype, which can be leveraged to enhance the healing process.

## Introduction

Following acute injury, tendons heal through a scar-mediated process, which leads to persistent functional deficits. A significant barrier to identify biological therapeutics to promote tendon regeneration is the limited understanding of the cellular and molecular programs, as well as cell-cell interactions governing healing. Scleraxis (*Scx*), a basic helix-loop-helix transcription factor, is a critical regulator of tendon formation^1, 2^, with loss of *Scx* resulting in developmental deficits in tendon. In addition, Scx-lineage (Scx^Lin^) cells play a critical role in tendon healing^3–5^, particularly in the formation of an organized cellular bridge between the injured tendon stubs^6^. However, defining the function of Scx^Lin^ cells and their associated molecular programs are complicated by the heterogeneity of this cell lineage. That is, the Scx^Lin^ cells encompass those cells that expressed *Scx* at some point during embryonic development, the progeny of these cells, as well as any cell that expresses *Scx* during postnatal growth, adult homeostasis, and in response to injury. We have previously shown that depletion of Scx^Lin^ cells in the adult flexor tendon improves the tendon healing and has only a modest impact on short-term tendon homeostasis^5^. Thus, we suggested that there are on-going ‘waves’ of cells being added to the Scx^Lin^ populations throughout the healing process, with distinct Scx^Lin^ subpopulations playing temporally- and spatially-dependent functions. While comprehensive definition of the functions of Scx^Lin^ cells is important, we are particularly interested in adult Scx^Lin^ cells (those cells that actively express *Scx* in the adult tendon during homeostasis) and their contribution to the myofibroblast fate during tendon healing. Myofibroblasts (identified as aSMA^+^) mediate healing via deposition, contraction and remodeling of new extracellular matrix (ECM)^7^. However, persistent or dysregulated myofibroblast function can promote a fibrotic healing response due to exuberant and sustained ECM deposition^7^. In many tissues the initiating step of myofibroblast differentiation is activation of quiescent, tissue-resident fibroblasts. We hypothesize that during fibrotic tendon healing, adult Scx^Lin^ resident tenocytes undergo fibroblast activation and subsequent myofibroblast differentiation in a tightly controlled temporal and spatially-dependent manner. In the current study, we have assessed the relationship between adult Scx^Lin^ cells and active *Scx* expression to elucidate the detailed molecular mechanisms underlying tenocyte-specific fibroblast activation and myofibroblast differentiation in tendon healing. Lineage tracing was used to define adult Scx^Lin^ spatial localization during activation and differentiation, and further used spatial transcriptomics to delineate transcriptional programs over time and begin to resolve crosstalk between heterogenous cell populations during tendon healing.

## Methods

### Animals Ethics

This study was carried out in strict accordance with the recommendations in the Guide for the Care and Use of Laboratory Animals of the National Institutes of Health. All animal procedures were approved by the University Committee on Animal Research (UCAR) at the University of Rochester.

### Mouse Models

Scx-Cre^ERT2^ and Scx-GFP mice were generously provided by Dr. Ronen Schweitzer. ROSA-Ai9 (#007909) mice were obtained from the Jackson Laboratory (Bar Harbor, ME, USA). ROSA-Ai9 mice fluoresce red following Cre-mediated recombination. Scx-Cre^ERT2^ mice were crossed to the ROSA-Ai9 strain to trace adult Scx-lineage cells (Scx^Ai9^), while Scx-GFP (Scx^GFP^) mice were used to track active Scx expression. For all experiments, Scx^Ai9^ animals received three 100mg/kg i.p. tamoxifen (Tmx) injections beginning seven days prior to flexor tendon repair surgery to trace and assess the fate of adult Scx^Ai9^ cells, while also ensuring that no subsequent labelling occurred during tendon healing by allowing for a washout period of four days. The only exception to this is in Figure 3C, in which mice received Tmx from days 10-12 post-surgery.

### Flexor Tendon Repair

At 10-12 weeks of age, male and female mice underwent complete transection and repair of the flexor digitorum longus (FDL) tendon in the hind paw as previously described^5, 8, 9^. Mice were injected prior to surgery with 15-20μg of sustained-release buprenorphine and then anesthetized with Ketamine (100mg/kg) and Xylazine (10mg/kg). Following cleaning and sterilization of the surgery area, the FDL tendon was transected at the myotendinous junction to reduce strain-induced rupture of the repair site and the skin was closed with a 5-0 suture. This myotendinous junction injury naturally heals throughout the first week of healing, re-introducing strain to the tendon. A small incision was then made on the posterior surface of the hind paw, the FDL tendon was located and completely transected. The tendon was repaired using an 8-0 suture and the skin was closed with a 5-0 suture.

### Histology and immunofluorescence

Hind paws were harvested (n=3-5 per time-point) from both uninjured Scx^Ai9^ mice, and Scx^Ai9^ mice at postoperative days 8, 10, 12, 14, 21, and 28 for paraffin sectioning, while tendons from Scx^Ai9^; Scx^GFP^ mice were harvested at 14, 21, and 28 days post-repair. Hind paws were fixed in 10% formalin for 72 hours at room temperature, decalcified in Webb-Jee EDTA for two weeks, processed and embedded in paraffin. Three-micron sagittal sections were cut, de-waxed, rehydrated, and probed with antibodies for tdTomato (1:500, AB8181, SICGEN), αSMA-CY3 (1:200, C6198, Sigma Life Sciences, St. Louis, MO, USA) or αSMA-FITC (1:500, F3777, Sigma Life Sciences), HSP47 (1:250, ab109117, Abcam), PCNA (1:100, ab29, Abcam, Cambridge, MA), FAP (1:500, ab53066, Abcam), VCAM1 (1:1000, ab134047, Abcam), GFP (1:500, ab290, Abcam). The following secondary antibodies were used: Donkey anti-mouse 488 (for PCNA) (1:200, #715-546-150, Jackson Immuno), Donkey anti-rabbit Rhodamine-Red-X (for FAP, Hsp47, VCAM-1, GFP) (1:200, #711-296-152, Jackson Immuno), Donkey anti-goat Rhodamine-Red-X (for tdTomato) (1:200, #705-296-147, Jackson Immuno), Donkey anti-goat 488 (for tdTomato) (1:200, #705-546-147, Jackson Immuno). Nuclei were counterstained with DAPI and imaging was performed using a VS120 Virtual Slide Microscope (Olympus, Waltham, MA, USA). For select slides following imaging, coverslips were gently removed in PBS from Scx^Ai9^ sections and samples subsequently stained with Masson’s Trichrome to visualize collagen content. Figures 2C and 3E were pseudo-colored using Image J (https://imagej.nih.gov/ij/) for ease of interpretation.

### Spatial Transcriptomics

Hindpaws were from Scx^Ai9^ mice harvested at 14 and 28 days post-surgery, and from un-injured Scx^Ai9^ mice, embedded in cryomatrix and snap-frozen in isopentane pre-cooled with liquid nitrogen. Note that hindpaws from two mice at 14 days post-surgery were harvested and served as biological replicates. Evaluation of batch effect for spatial transcriptomics analysis were also conducted on these two D14 samples (Supplementary Figure 1). Ten-micron frozen sections were cut through the sagittal plane of the tendon and mounted on a spatially barcoded array slide (Visium Spatial Gene Expression, 10X Genomics). Sections were fixed and stained with H&E for digital imaging, permeabilized and then a reverse transcription mix was added to each section. The bound cDNA was then released from the slide, collected and prepared for RNA sequencing. Libraries were sequenced on the Illumina Nextseq platform (Illumina Inc, San Diego CA) using paired-end sequencing. Sequence data was then overlaid on to the H&E stained digital image based on spatial orientation of the barcodes. The 10X gene-expression data were first processed using CellRanger (v.3.0.2, 10X Genomics). Normalized feature-barcode matrices was then used for downstream analysis.

### Spatial RNA-seq data processing, integration, and visualization

Spatial RNA-seq datasets of the samples from different timepoints were processed using Seurat V4.0 R package^10^. Each dataset was normalized based on regularized negative binomial models with the *SCTransform* function. Dimensionality reduction of datasets were performed by *RunPCA* function with 50 principal components (npcs = 50) as an argument. We next used the *FindNeighbors* function to compute the shared nearest-neighbors (SNN) for a given dataset with parameter k = 20. Clusters of the cells were identified based on SNN modularity optimization with *FindClusters* function. *RunUMAP* function was further used to perform Uniform Manifold Approximation and Projection (UMAP) dimensional reduction^11^. Cell clusters were visualized on reduced UMAP dimensions using *DimPlot* function and on anatomical regions of tissue sections using the *SpatialDimPlot* function. To identify molecular features that correlate with spatial location within a tissue, we used the *FindSpatiallyVariableFeatures* function to choose the top 1000 highly variable genes from the dataset using the “markvariogram” selection method. Expression of genes of interest was visualized using the *SpatialFeaturePlot* function.

### Batch effect evaluation for D14 Spatial RNA-seq samples

Clusters including scar and tendon tissues were subsetted from both D14 samples (biological replicates) in Seurat v4.0. Subsetted clusters were log-normalized with scale factor 1000 and obtained 2000 highly variable genes using the *NormalizeData* and *FindVariableFeatures* functions, sequentially. The batches of the samples were then integrated and corrected with an anchor-based approach implemented in Seurat using *FindIntegrationAnchors* and *IntegrateData* functions. Visualization of corrected batches (Supplemental Figure 2) demonstrated similar cell numbers and distribution of cell clusters in UMAP space between two samples, suggesting our injury outcomes and sequencing results are highly reproducible.

### Integration of multiple Spatial RNA-seq datasets

To investigate dynamic changes in transcriptomic profiles of cell populations within scar and tendon tissues over time, we integrated spatial RNA-seq datasets from various timepoints prior and post-surgery. First, potential tendon and scar tissues were subsetted from each individual dataset (i.e. cluster 4 from the uninjured sample, clusters 0, 1, and 8 from D14 sample 1, clusters 1 and 3 from D14 sample 2, and clusters 2, 6, and 7 from D28). Subsetted datasets containing clusters of interest were log-normalized and scaled as previously described. Highly variable genes in the each subsetted datasets were identified using the *FindVariableFeatures* function. We then used the *FindIntegrationAnchors* and *IntegrateData* functions to align multiple datasets. After dataset integration, dimensionality reduction on the data was performed by computing the significant principal components on 2000 highly variable genes. Unsupervised clustering was done with the *FindClusters* function in Seurat with resolution argument set to 0.5, and clusters visualized with a UMAP plot. Differentially expressed genes (DEGs) among each cell cluster were determined using *FindAllMarkers* function. DEGs expressed in at least 25% cells within the cluster and with a fold change of more than 0.25 in log scale were considered to be marker genes of the cluster. To determine the biological functions of the marker genes from a given cluster, we performed Gene Ontology (GO) enrichment analysis using DAVID Gene Functional Classification Tool (http://david.abcc.ncifcrf.gov; version 6.8)^12^ with the top 100 genes of each cluster. By comparing these unique biological GO terms with current literature, we were able to annotate cell clusters. Additionally, the top 10 enriched GO terms from Biological Process category with associated p values were visualized using GraphPad Prism (version 9.0; GraphPad Software). To identify DEGs for the cells of the same type but across different timepoints, we used the *FindAllMarkers* function. DEGs with adjusted p value > 0.05 and log2 fold change > 2 were visualized by heatmaps using the ComplexHeatmap R package^13^.

### Pseudotemporal ordering and lineage trajectories

We used Monocle3 R package to reconstruct differentiation trajectories of tenocyte-fibroblastic scar tissue cells by computing and ordering the sequence of gene expression changes of the cells collected from different time points^14, 15^. First, we removed the Mmp3/Mmp9 enriched bridging tissue cluster from the integrated spatial-RNA-seq dataset, as we believe this cell population represents inflammatory macrophages due to its co-expression of *Cd11b* (*Itgam*), *Cd45* (*Ptprc*), F4/80 (*Adgre1*) as well as high expression levels of *S100a8, S100a9, IL1b*, and several Mmp genes (Supplementary Figure 3). Next, the spatial RNA-seq dataset created in Seurat was converted into a Monocle3 object using SeuratWrappers R package. The cells were then reclustered by *cluster_cells* function with a resolution of 0.01 in Monocle3. We used the *learn_graph* and *order_cells* functions to identify potential differentiation trajectories in our integrated spatial-seq dataset with the non-reactive/ native tendon-like cell cluster as the root of lineage trajectories.

To determine sets of genes (i.e., gene modules) governing the differentiation of non-reactive/ native tendon-like cells into either peripheral fibroblastic cells or highly reactive remodeling tendon cells during wound healing, we used the *graph_test* and *find_gene_modules* functions with resolution of 0.001 and k = 13 as arguments. Gene modules were then organized by their similarity using hierarchical cluster analysis. With this approach, we identified that gene module 2 contains genes controlling cell fate decision of non-reactive/ native tendon-like cells into highly reactive remodeling tendon cells, while genes in module 5 regulate the differentiation of non-reactive/native tendon-like cells into peripheral fibroblastic cells (Supplemental Figure 7). Additionally, GO term analysis was performed on gene module 2 and module 5 for their associated biological functions.

### Transcription factor (TF) binding motif and protein-protein network analysis

To obtain TF scores and TF binding motifs of the genes within modules that control lineage specification of non-reactive/native tendon-like cells during wound healing, we used genes from module 2 and module 5, respectively, as input gene lists to RcisTarget R package^16^ with default parameters and *mm9-tss-centered-10kb-7species.mc9nr.feather* as the database. Furthermore, gene lists from module 2 and 5 were submitted to Transcriptional Regulatory Relationships Unrevealed by Sentence-based Text mining (TRRUST; available at https://www.grnpedia.org/trrust/), a manually curated database of human and mouse transcriptional regulatory networks, to compare and combine the results obtained from RcisTarget analysis. For each gene module, top 3 TFs with corresponding binding motif were selected and visualized.

To determine protein-protein network within the gene module that regulates cell fate decisions of non-reactive/ native tendon-like cells, we input gene lists into STRING v11.0, a database of known and predicted protein-protein interactions covering 24,584,628 proteins (https://string-db.org/cgi/input?sessionId=bYLAXihSg3i9&input_page_show_search=on)^17^. We used the highest confidence (0.900) for minimum required interaction score and removed disconnected nodes in the network. The constructed network was further clustered using k-means clustering with maximum of 3 clusters as criteria (Supplemental Figures 8 & 9).

### Cell-Cell interactome analysis

To identify cell-cell crosstalk between inflammatory macrophage cells and highly reactive remodeling tendon cells, we used CellPhoneDB (v.2.1.7) package^18^, a publicly available repository of curated receptors, ligands and their interactions, in a Python v3.8 environment. As CellPhoneDB currently only supports cell-cell interactions identified in *Homo Sapiens*, we first converted enriched genes in each cell cluster into human homologue of the mouse genes using *Esembl* (http://useast.ensembl.org/biomart/martview/4bbd665c11e2de67f9aa5af2d58308ab)^19^. Converted genes were then used as input with stringent criteria (i.e., 50% of cells expressing the ligand/receptor and interactions with p value < 1e-20) were considered significant and the result was visualized using the *dot_plot* function embedded in CellPhoneDB package.

## Results

### Adult Scx^Ai9^ cells break quiescence following injury and take on an ‘activated’ phenotype during early healing

While tenocytes are largely quiescent prior to injury^20^, there is an active involvement of proliferative cells, including those from the tendon periphery during healing^21, 22^. To assess *in vivo* proliferative capability of Scx^Ai9^ tendon cells during flexor tendon healing (Fig. 1A), Scx^Ai9^ tendons were stained for proliferating cell nuclear antigen (PCNA) at days 8, 10, 12, 14, 21, and 28 post-repair. While there are PCNA+ Scx^Ai9^ cells in both the tendon stub and bridging scar tissue at days 8 and 10 post-repair, proliferation of Scx^Ai9^ cells appeared to peak between days 12 and 14 at both the stub and bridging regions. While many Scx^Ai9^ cells within the stub remained PCNA+ at day 21 post-repair, bridging Scx^Ai9^ cells were almost exclusively PCNA-. By day 28 post-repair, only a few Scx^Ai9^ cells in the stub were PCNA+ and few of bridging Scx^Ai9^ cells, if any, appeared to express PCNA marker (Fig.1B). Therefore, bridging Scx^Ai9^ cells may be derived from both newly proliferative cells that emanate from the stub and proliferative Scx^Ai9^ cells that have already migrated into the bridge from peripheral tendon tissues.

**Figure 1.**
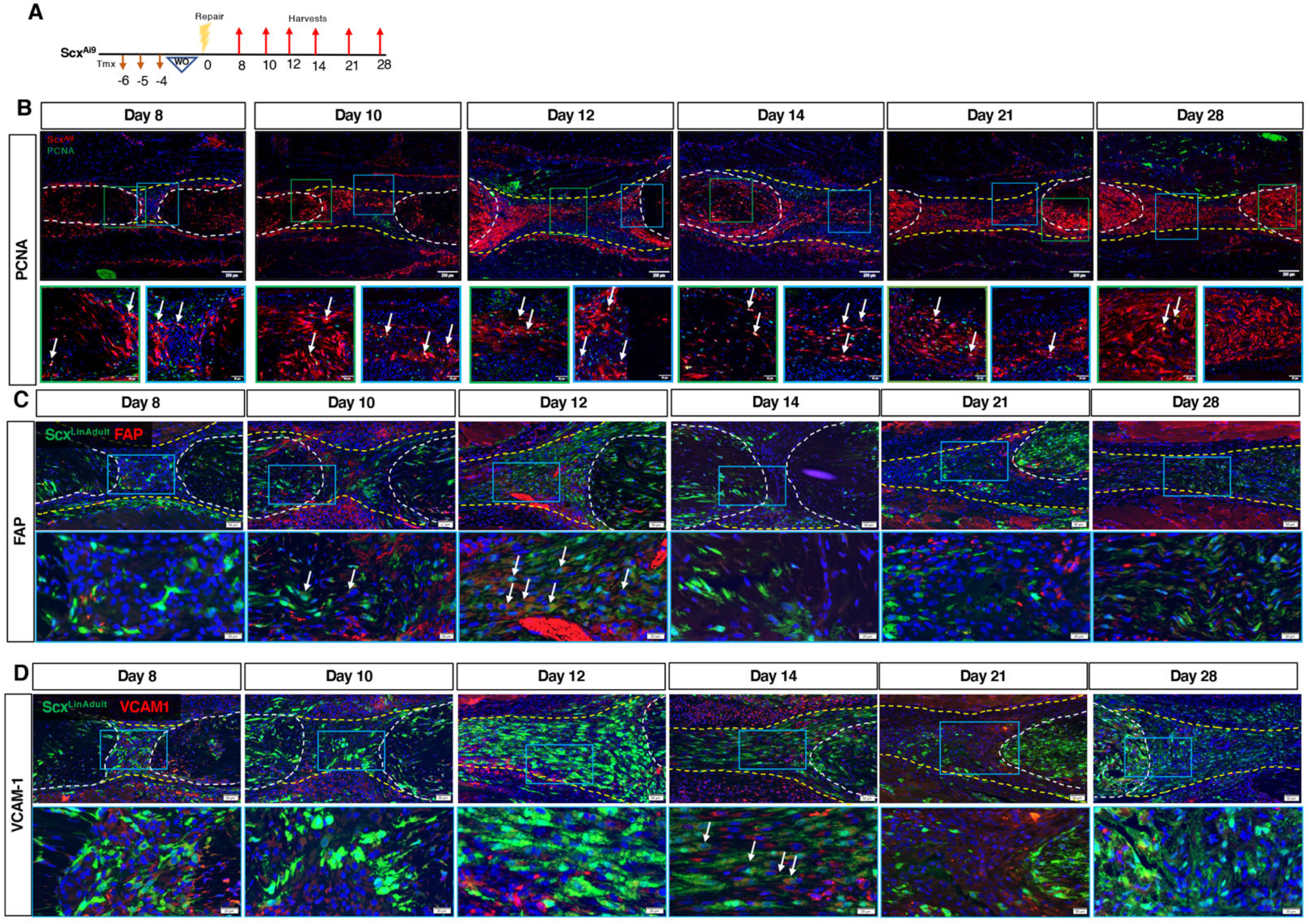
Scx^Ai9^ cells residing in both the tendon stubs and bridging tissue proliferate through day 21 post-repair. (A) Scx^Ai9^ mice were injected with Tmx for three days, followed by a four day washout (WO) period prior to tendon repair surgery, and tendon were harvested at 8, 10, 12, 14, 21, and 28 days post-repair. (B) Co-immunofluorescence of Scx^Ai9^ (red) and PCNA (green), (C) Scx^Ai9^ (green) and FAP (red), (D) Scx^Ai9^ (green) and VCAM-1 (red) between 8-28 days post-repair. Nuclei are stained with DAPI. Tendon is outlined by white dotted lines and scar tissue by yellow dotted lines. Green and blue boxes indicate location of higher magnification images Examples of co-localization indicated by white arrows. N=3-4 per timepoint.

To determine spatial-temporal patterns of fibroblast activation of adult Scx^Ai9^ cells during tendon healing, fibroblast activation protein (FAP) and vascular cell adhesion protein 1 (VCAM-1) were evaluated. At D8 there was minimal expression of FAP, and by D10 there were a few FAP+/Scx^Ai9^+ cells (white arrows, Fig. 1C), but most FAP expression was adjacent to Scx^Ai9^ cells (Fig. 1C). By D12, most Scx^Ai9^ cells in the bridging tissue were FAP+; however, by D14, FAP expression had dramatically decreased in both the bridging tissue and tendon stubs, with only a few FAP+ cells remaining in the tendon stubs. These low levels of FAP expression persisted through D28. At D8, several Scx^Ai9^+ cells co-expressed VCAM-1, while some VCAM-1 positive cells were Scx^Ai9^ negative (Fig. 1D). The VCAM-1 expression pattern increases through D14, at which time many Scx^Ai9^+ cells in the bridging tissue are VCAM-1 positive. A substantial decrease in overall VCAM-1 expression, and VCAM-1+/Scx^Ai9^+ cells are observed at D21 and D28. Collectively, these data demonstrate that Scx^Ai9^ cells proliferate throughout the healing process, and undergo transient fibroblast activation, with these processes proceeding in a spatially-dependent manner.

### The interior organized ECM bridge is formed by adult Scx^Ai9^ cells

Fibroblast activation is associated with both matrix production and myofibroblast differentiation. While we have previously demonstrated the adult Scx^Ai9^ cells form a highly aligned cellular bridge between the native tendon ends ^6^, it is unclear whether these cells elaborate this new ECM or simply use an existing ECM scaffold as a bridge during tendon repair. To address this, we examined both Scx^Ai9^ cell localization and ECM deposition in the same samples using sequential immunofluorescence and Masson’s trichrome staining. As we have previously demonstrated a lack of both Scx^Ai9^ cells, and new collagen ECM between the native tendon ends at D7^6^, we analyzed samples starting at D8 post-surgery (Fig. 2A) in the current study. At D8 a few Scx^Ai9^ cells have egressed from the tendon stubs concomitant with modest collagen fiber presence (Fig. 2B). At D10, a substantial increase in the extent of both Scx^Ai9^ and collagen ECM are observed, with strong overlap between the cells and ECM, and this tandem elaboration of the cell/ECM bridge continues through D12 (Fig. 2B). These data demonstrate concomitant development of the Scx^Ai9^ cellular bridge and the collagen ECM bridge, suggesting a key role for Scx^Ai9^ cells in elaboration of this new, organized matrix. Interestingly, there is a clear demarcation between Scx^Ai9^+ cells and Scx^Ai9^-cells at periphery of the new ECM bridge (yellow arrows, Fig. 2B), suggesting the presence of discrete molecular programs and cell populations in this area. We then examined expression of the collagen chaperone protein, Hsp47, which shows abundant colocalization of Hsp47 in Scx^Ai9^ cells as the new ECM is developed (Fig. 2C), further supporting Scx^Ai9^ cells as the primary producers of this new ECM bridge.

**Figure 2.**
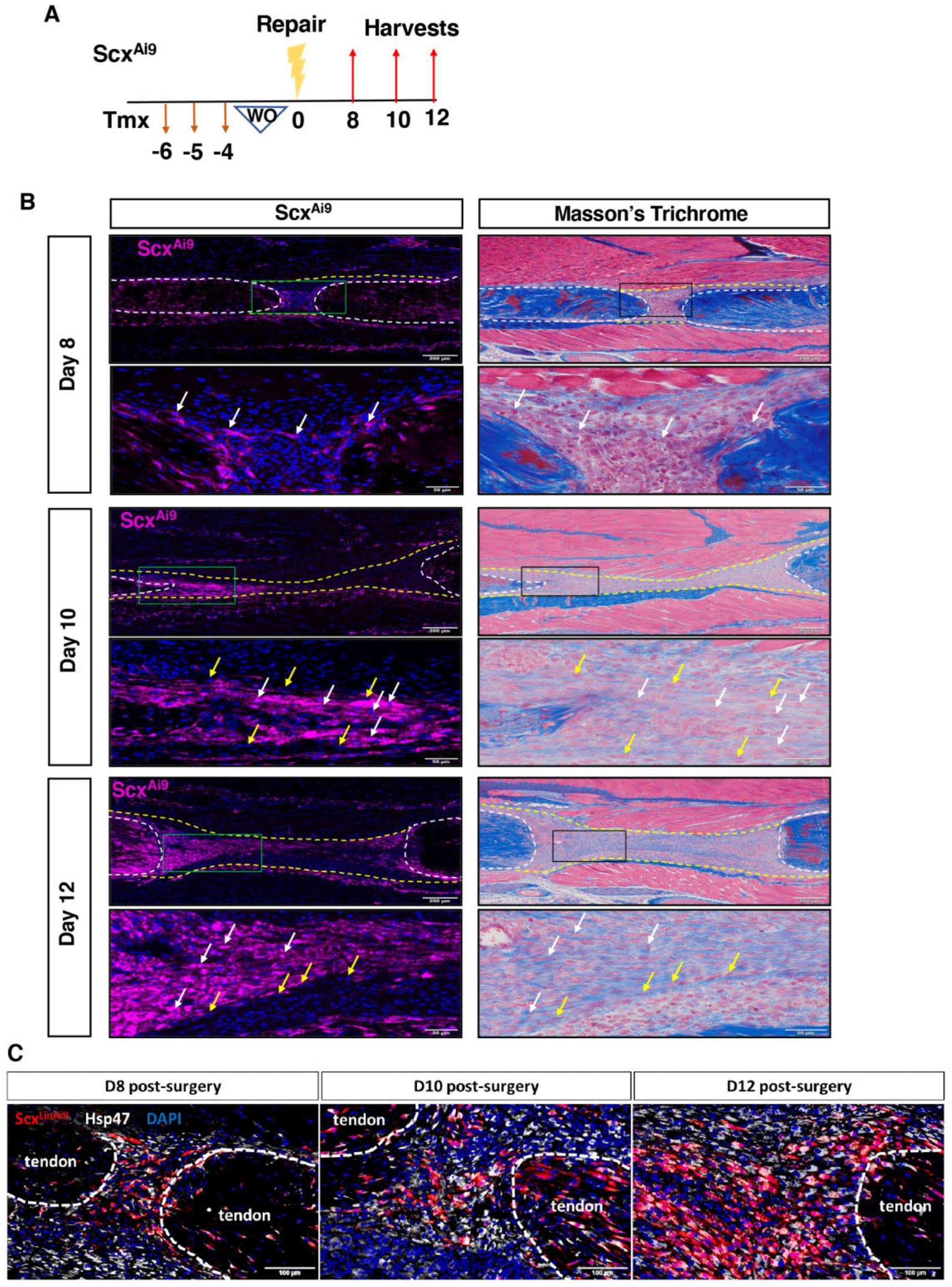
Bridging Scx^Ai9^ cells and bridging collagen matrix both actively form by 10 days post-repair. (A) Scx^Ai9^ mice were injected with Tmx for three days, followed by a four day washout (WO) period prior to repair surgery. Healing tendons were harvested at days 8, 10, and 12 post-repair. (B) Immunofluorescence of Scx^Ai9^ (pink) at days 8, 10, and 12 post-repair. Nuclei stained with DAPI. Coverslips were removed and samples were then stained with Masson’s Trichrome to analyze localization of organized collagen deposition at the repair site. Tendon is outlined by white dotted lines and scar tissue by yellow dotted lines. Green and black boxes indicate location of higher magnification images. White arrows used to indicate Scx^Ai9^ cells and yellow arrows used to indicate non-Scx^Ai9^ cells between fluorescent and masson’s trichrome images. (C) Co-immunoflourescence for Scx^Ai9^ (red) and Hsp47 (white) between D8-12 post-repair. Tendon stubs are outlined in white. N=3-4 per timepoint.

### Adult Scx^Ai9^ cells undergo transient myofibroblast differentiation, with an additional population of Scx^Lin^ cells contributing to the myofibroblast fate over time

We have previously shown that adult Scx^Lin^ cells are not the primary contributor to myofibroblast fate at D14^6^; however, based on the transient ‘activated’ state of Scx^Ai9^ cells, and the ECM elaboration phenotype, we examined subsequent myofibroblast fate more comprehensively (Fig. 3A). At D8 and D10, minimal alpha smooth muscle actin (aSMA)+ myofibroblasts were observed. However, by D12 an abundant aSMA+ population was observed, with many of these cells derived from Scx^Ai9^+ cells. However, consistent with our previous data, at D14 aSMA+ Scx^Ai9^-derived myofibroblasts were primarily restricted to the tendon stubs and not in the bridging granulation tissue (Fig. 3B), with this expression pattern continuing through D28, indicating spatially-dependent myofibroblast differentiation. Interestingly, these data suggest that not all adult Scx^Ai9^ cells that undergo fibroblast activation continue to a mature myofibroblast phenotype, and that contribution to the myofibroblast fate is both time and location dependent.

**Figure 3.**
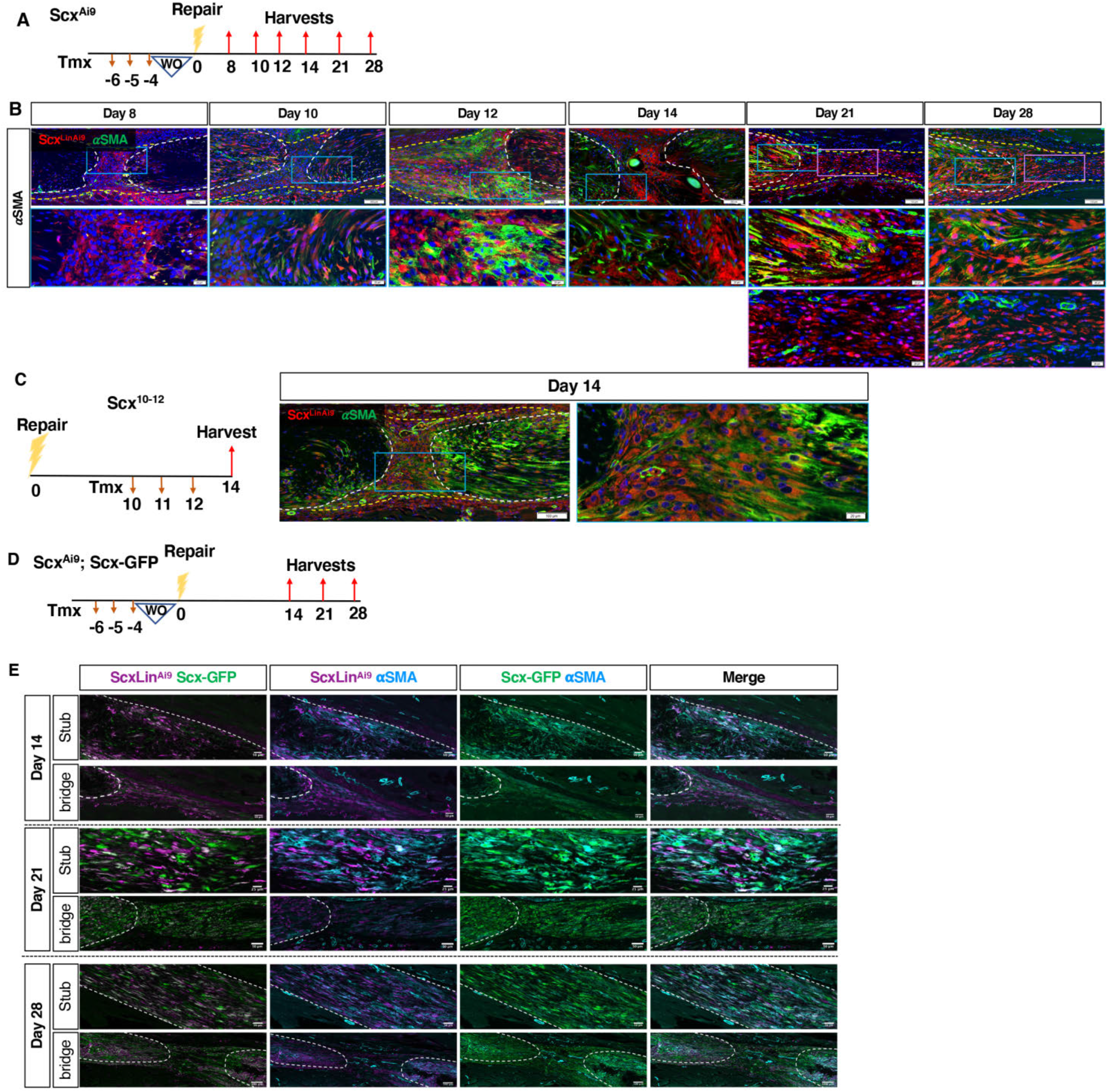
Bridging Scx^Ai9^ cells do not differentiate into αSMA+ myofibroblasts. (A) Scx^Ai9^ mice were injected with Tmx for three days, followed by a four day washout (WO) period prior to tendon repair surgery. Tendons were harvested between 8-28 days post-repair. (B) Co-immunofluorescence between Scx^Ai9^ (red) and αSMA+ myofibroblasts (green) between 8-28 days post-repair. Blue and purple boxes indicate locations of higher magnification images. (C) Scx^Ai9^ mice were injected with Tmx on days 10-12 post-repair and harvested on day 14 to label cells actively expressing *Scx* during healing (Scx^10–12^). Co-immunofluorescence between Scx^10–12^ (red) and αSMA+ myofibroblasts (green). Blue box indicates location of higher magnification Examples of co-localization indicated by white arrows. Nuclei stained with DAPI. (E) Scx^Ai9^; Scx^GFP^ mice were injected with Tmx for three days, followed by a four day washout (WO) period prior to tendon repair surgery. Tendons were harvested between 14-28 days post-repair, and stained for Scx^Ai9^ (purple), Scx^GFP^ (green), and αSMA+ myofibroblasts (blue).Tendon is outlined by white dotted lines and scar tissue by yellow dotted lines. N=3-5 per timepoint.

It has previously been demonstrated that approximately 65% of αSMA-lineage cells express *Scx* at day 14 post-injury in patellar tendon ^3^. To confirm that actively expressing αSMA+ myofibroblasts express *Scx* postinjury, Scx-Cre^ERT2^; ROSA-Ai9^F/F^ mice were injected with Tmx on days 10-12 post-repair to label cells expressing *Scx* near the time of a day 14 harvest (Scx^10–12^) (Fig. 3C). Essentially all αSMA+ myofibroblasts express *Scx* by day 14 post-flexor tendon repair (Fig. 3C), suggesting that *Scx* expressing myofibroblasts in the bridging region are not derived from Scx^Ai9^ cells that are labelled in the adult tendon prior to injury, but instead from a separate subpopulation of Scx^Lin^ cells that actively express Scx.

Based on these data, and our previous work demonstrating that the broad Scx^Lin^ cell pool (Scx-Cre) contributes to a persistent myofibroblast population^23^, we wanted to better define this population. Given that there are likely on-going ‘additions’ to this overall Scx^Lin^ pool during healing^5^ with cells turning on *Scx* expression at different times post-injury, we aimed to gain a more comprehensive understanding of the relationship between active *Scx* expression, adult Scx^Ai9^ cells and myofibroblast fate. To investigate this, we generated Scx^Ai9^; Scx^GFP^ dual-reporter mice and traced these cells’ myofibroblast fate during tendon healing (Fig. 3D & E).

At D14 there were populations of Scx^Ai9^+/Scx^GFP^-and Scx^Ai9^-/Scx^GFP^+ cells in both the tendon stubs and the bridging tissue, but Scx^Ai9^+/Scx^GFP^+ seemed to be the predominant cell type in both the tendon stubs and the bridging tissue. Interestingly, this Scx^Ai9^+/Scx^GFP^+ population was substantially decreased in the tendon stubs and bridge at D21, with a predominance of Scx^GFP^+ cells in the tendon bridge. In the stubs, both Scx^Ai9^+/Scx^GFP^-and Scx^Ai9^-/Scx^GFP^+ populations were present; however, Scx^Ai9^-/Scx^GFP^+ was the primary population. By D28, there was a substantial shift in populations in the stubs with a predominance of Scx^Ai9^+/Scx^GFP^+, and a few Scx^Ai9^-/ScxGFP+. Interestingly, most Scx^Ai9^+ were also Scx^GFP^+. The large presence of Scx^GFP^+ continued in the bridging tissue at D21, though there was a progressive decrease in Scx^GFP^+ and proximity to the tendon stubs increased (Fig 3E). These data are consistent with our previous hypothesis^5^ that the contribution of adult Scx^Ai9^ cells are overwhelmed during early healing by an influx or expansion of a Scx+ population, followed by resolution of this population, and predominance of adult Scx^Ai9^ cells (particularly in the tendon stubs) during later healing.

In terms of myofibroblast fate, consistent with our previous data^6^ (Fig. 3B), Scx^Ai9^+ cells primarily contribute to the myofibroblast fate in the stubs between D14-28. In addition, at D14, most αSMA+ myofibroblasts in the tendon stubs are Scx^GFP^+, with minimal αSMA+ cells in the tendon bridge, with this pattern continuing at D21 and D28. Interestingly, Scx^GFP^ expression was retained in nearly all αSMA+ myofibroblasts in the tendon stubs, with these cells being predominantly derived from Scx^Ai9^ at D14 and D21, and almost exclusively from Scx^Ai9^ at D28 (Fig. 3E). Collectively, these data demonstrate both time and location-dependent contributions of adult Scx^Ai9^ and Scx^GFP^ cells to myofibroblast fate.

### Defining the spatially mapped cellular populations in fibrotic tendon healing

The clear spatial dependence of the adult Scx^Ai9^ contribution to ECM organization and persistent myofibroblast populations suggests that there are likely discrete cellular populations and molecular programs underlying these fate-function dynamics. Therefore, we utilized spatial transcriptomics to identify the spatial distribution of heterogenous cellular populations and their associated gene expression profiles in tendon healing.

An initial unsupervised clustering of each spatial map of the hind paw identified 6 distinct clusters in the uninjured hind paw, 10 clusters in D14 sample 1, 7 clusters in D14 sample 2, and 8 clusters in D28, respectively (Figure 4A, B, E, F; Supplemental Figure 1). Furthermore, no apparent batch effects were observed in two independent D14 samples (Supplemental Figure 2). From there, specific clusters that encompassed our area of interest (tendon and scar tissue) were selected for subsequent analysis. For uninjured tendon, Cluster 4, which overlapped with tendon tissue was selected for subsequent analysis and integration with post-operative timepoint (Supplemental Figure 1). At D14, clusters 0, 1, and 8 from the unsupervised clustering of the hind paw were selected due to their overlap with native tendon and the repair site (Fig. 4C). Re-clustering three clusters at D14 results into five sub-clusters (0-5) were identified, with cluster 1 primarily in the bridging granulation tissue between the tendon stubs and in the reactive/remodeling tendon stubs immediately adjacent to the repair site. Matrix metalloproteinases and tissue inhibitors of MMPs (*Mmp13, Mmp9, Timp1*), as well as myofibroblast marker *Acta2* (aSMA) were upregulated in cluster 1, implying active remodeling/matrix production at this site (Fig 4D). Clusters 2 and 3 both correspond to areas of native tendon immediately adjacent to the repair site (Fig. 4C), indicating some degree of spatio-molecular heterogeneity. While both these clusters share high expression of *Col1a1*, as expected, Cluster 2, which had high *tdTomato* (Scx^Ai9^) expression, is denoted by upregulation of *Fmod, Coch* and *Lox*, while cluster 3 is enriched *Fn1, Bgn*, and *Tnmd*, suggesting complementary ECM production/organization roles for each tendon end. Cluster 0 appears to surround the injury site, and expression of *Fbn1, Tppp3*, and *Postn* is highly upregulated, indicating that this cluster may represent a stem-cell-like pool ^24^. Finally, high expression of *Cd6* and *Csf1r* in cluster 4 indicates that this group corresponds to an infiltrating immune cell population (Figure 4D).

**Figure 4.**
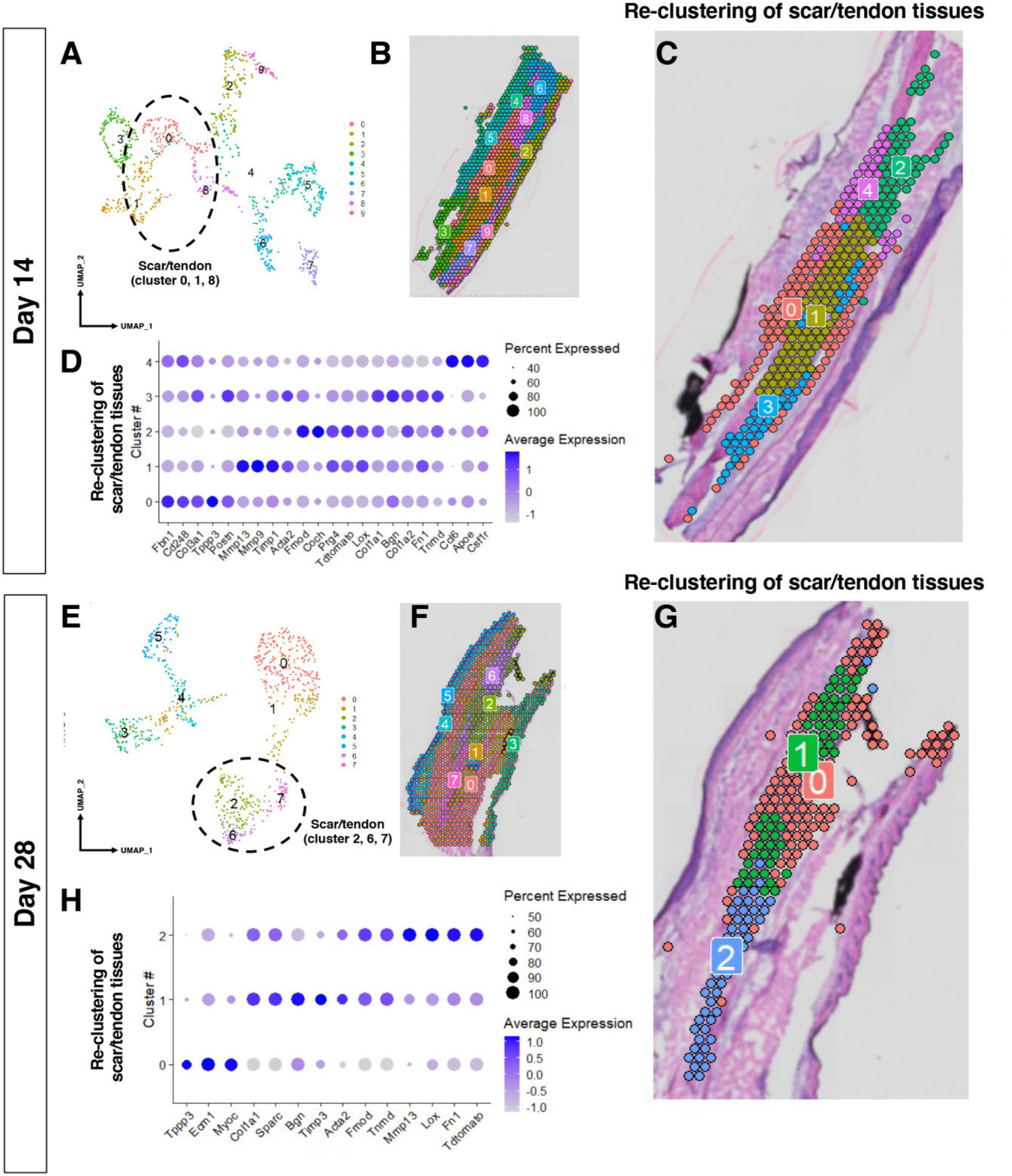
Spatial transcriptomics analysis of tendon healing. (A) Unsupervised clustering of the transcriptome of D14 post-repair. (B) Mapping of the unsupervised clusters on to the histological section at D14. (C) Mapping of the reclustering of the selected clusters that overlap with tendon and scar tissue on the slide at D14. (D) Dot plot of the genes that are enriched within each of the clusters at D14. (E) Unsupervised clustering of the transcriptome of D28 post-repair. (F) Mapping of the unsupervised clusters on to the histological section at D28. (F) Mapping of the reclustering of the selected clusters that overlap with tendon and scar tissue on the slide at D28. (H) Dot plot of the genes that are enriched within each of the clusters at D28.

At D28, clusters 2, 6, and 7 from the unbiased analysis underwent an additional clustering step, resulting in 3 subclusters (Clusters 0, 1, 2) (Fig. 4G). The resulting clusters 1 and 2 represent two complementary molecular programs for collagen organization at the site of injury, enriched in *Col1a1/Sparc/Bgn* and *Tnmd/Mmp13/Lox* respectively (Fig. 4H). Cluster 0 appears to closely resemble cluster 4 in the D14 reclustered sample, with the same localization surrounding the repair site, and enrichment of *Tppp3, Ecm1*, and *Myoc* (Fig. 4H).

### Integration of multiple spatial RNA-seq datasets and elucidation of dynamic changes in transcriptomic profiles of cellular subpopulations overtime

Integrating the selected clusters (i.e., tendon and scar tissue) from each timepoint allows us to identify heterogenous subpopulations at higher resolution and determine the dynamic changes in transcriptomic profiles overtime. After datasets integration, 5 conserved cell clusters over the course of tendon healing were identified (Fig. 5A). Examination of DEGs for each cluster coupling with their spatial destructions allow approximate annotation for these cellular populations. Cluster 0 corresponds to a peripheral, fibroblastic tissue capsule, enriched in *Pi16, CD248* and *Col14a1* (Fig. 5A & B). GO term analysis reveals that this population is involved in collagen fibril organization and cell adhesion, reinforcing this annotation, and interestingly this cluster also appears to be heavily involved in the complement system and innate immune response (Supplemental Figure 4). Spatial location of the cluster 1, enriched in *Coch, Chad*, and *Car3*, most closely approximates native tendon-like tissue, and GO term analysis indicates upregulation of contractile genes (Supplemental Figure 4). Further away from the injury site, cluster 2 represents a less reactive state, identified by *C1qtnf3, Tnmd*, and *Col1a1* and others primarily involved in ECM organization pathways (Supplemental Figure 5). Cluster 3 marks the bridging tissue, and corresponds to a more inflammatory program, enriched in *Acp5, Cd11b* (*Itgam*), *Cd45* (*Ptprc*), F4/80 (*Adgre1*) as well as high expression levels of *S100a8, S100a9, IL1b*, and several Mmp genes (Supplemental Figure 3). Pathway analysis confirms the role of this cluster in collagen catabolism and immune response, suggesting that infiltrating macrophages may make up part of this group (Supplemental Figure 3 & 5). Finally, cluster 4 identifies the highly reactive, actively remodeling tendon immediately adjacent to the repair site, with increased expression of *Spp1, Mmp13*, and *Timp1* and upregulation of pathways involved in collagen catabolism, ECM organization, and wound healing (Fig. 5B, Supplemental Figure 6).

**Figure 5.**
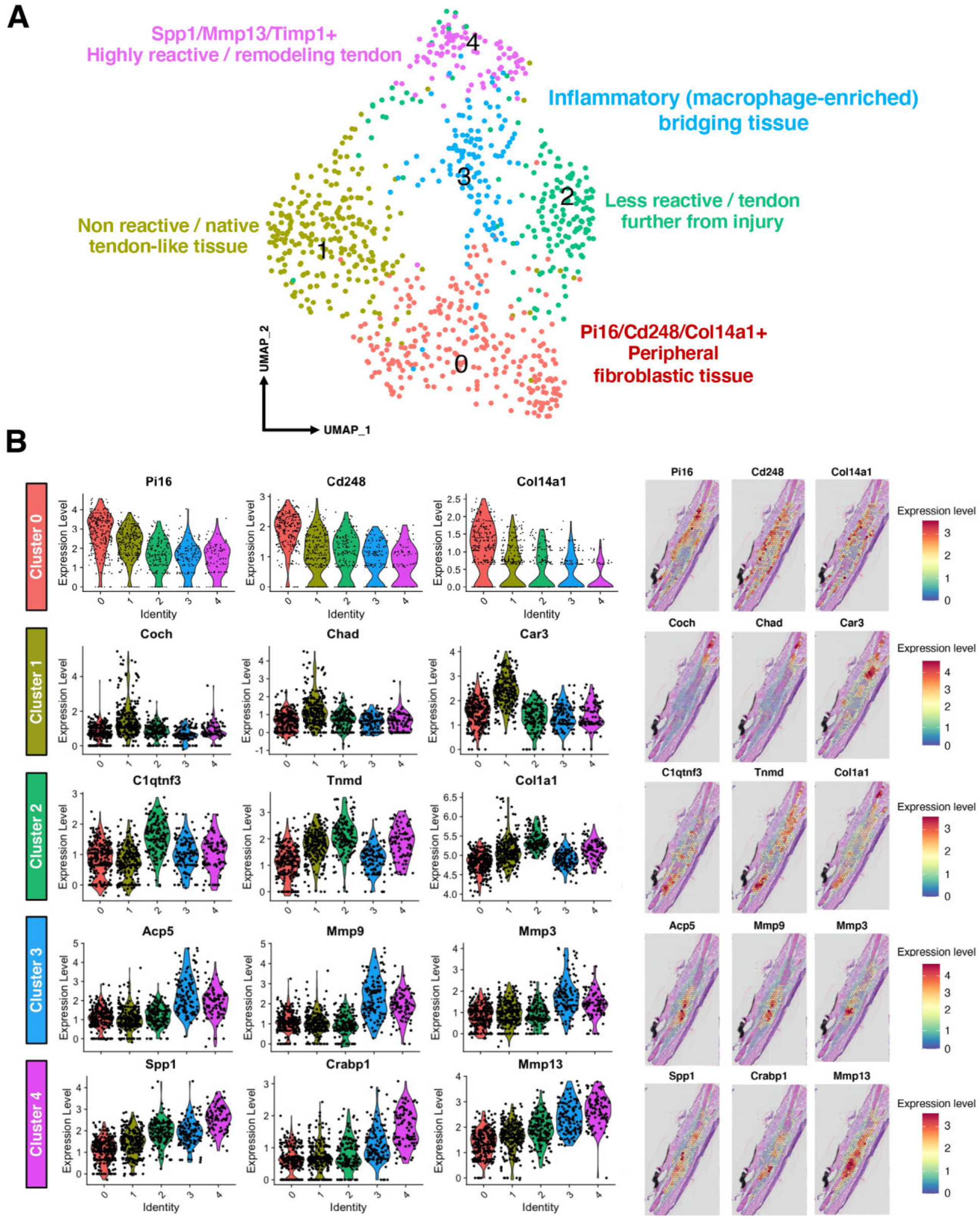
Integration of tendon and scar subclusters defines transcriptome dynamics over time. (A) UMAP of integrated data from subclusters overlapping with tendon and scar from uninjured, D14 and D28 identifies five subpopulations. (B) Violin plots and spatial mapping of highly expressed genes within each integrated cluster.

### Identification of fibroblastic differentiation trajectories and key transcriptional regulators

To better understand the spatially-dependent differentiation trajectories of fibroblastic cells over the course of healing, pseudotime analysis was conducted including all clusters except inflammatory macrophage (cluster 3) due to their potential myeloid origin (Supplemental Figure 3), using the non-reactive/native tendon-like cell cluster (cluster 1) as the root of lineage trajectories. A lineage bifurcation was observed for non-reactive/ native tendon-like cells (cluster 1) as they may differentiate either into highly reactive remodeling tendon cells (cluster 4) or peripheral fibroblastic cells (cluster 0) during tendon healing (Fig. 6A & B, Supplemental Figure 7). We next analyzed the genes that are co-regulated around trajectory branch point to further obtain insights into the genetic profiles governing cell fate decisions. Trajectory branch analysis using Monocle3 identified 14 gene modules that may contribute cell fate specification of non-reactive/ native tendon-like cells (cluster 1) (Supplemental Figure 7): module 2 may contain genes controlling cell fate decision of non-reactive/ native tendon-like cells (cluster 1) into highly reactive remodeling tendon cells (cluster 4), while genes in module 5 may be responsible for driving non-reactive/ native tendon-like cells (cluster 1) into peripheral fibroblastic cells (cluster 0).

**Figure 6.**
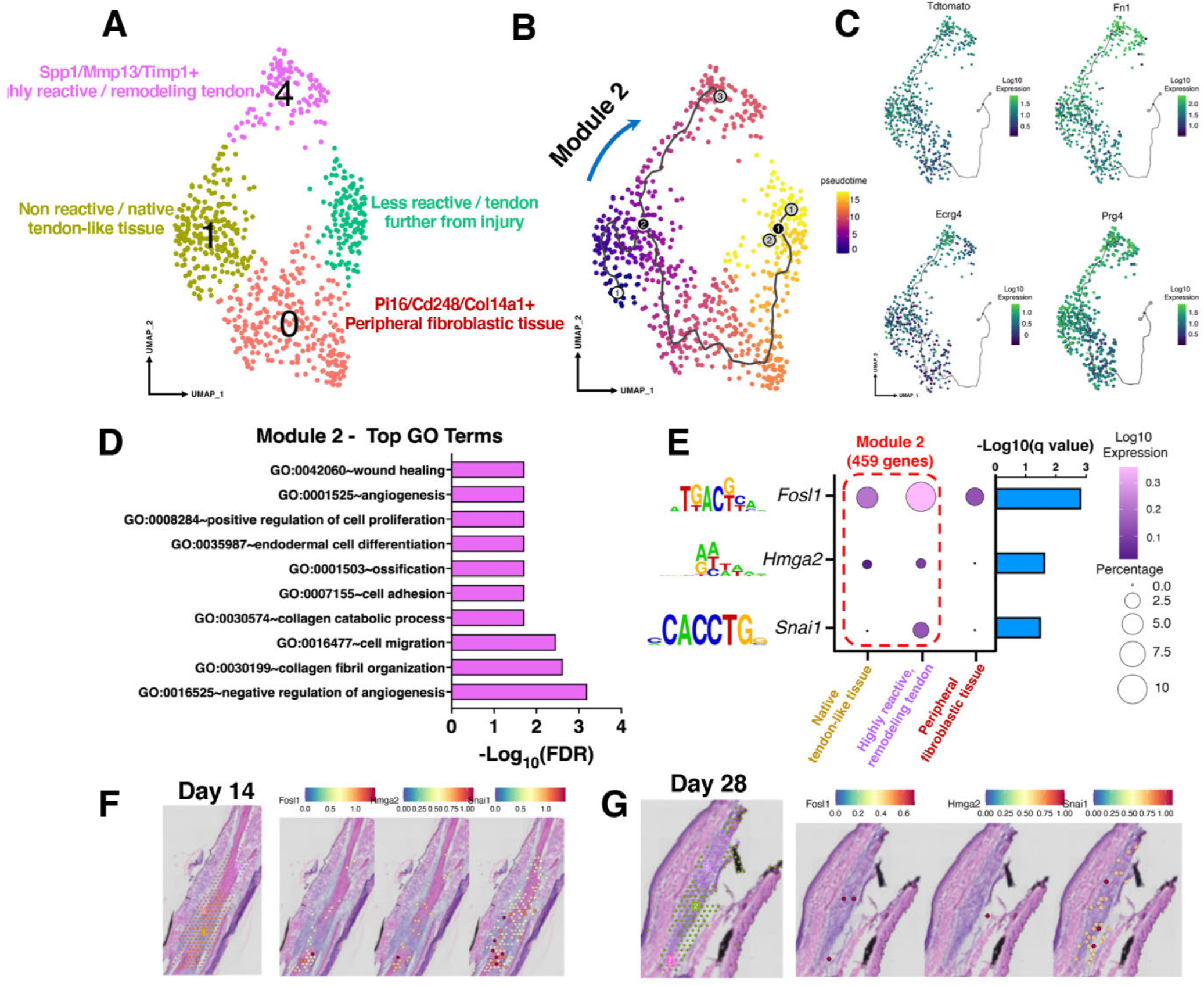
Pseudotime trajectory analysis from basal tendon to reactive tissue. (A) UMAP of the clusters included in pseuodotime analysis (Macrophage cluster was excluded). (B) Pseudotime analysis demonstrating the location of module 2 which was used for subsequent analysis. (C) Identification of genes that define the trajectory in module 2, including *tdTomato, Fn1, Ecrg4*, and *Prg4*. (D) Identification of GO terms enriched in module 2 trajectory. (E) TF binding motif analysis using the RcisTarget R package predicts transcription factors *Fosl1, Hmga2* and *Snai1* as regulators of the trajectory in module 2. (F & G) Spatial mapping of the subclusters included in pseudotime analysis and the predicted transcription factors regulating this trajectory at D14 (F) and D28 (G).

The differentiation trajectory of module 2 demonstrated increased expression levels of *Fn1, Ecrg4, tdTomato*, and *Prg4* when native tendon-like cells (cluster 1) gradually differentiated into highly reactive remodeling tendon cells (cluster 4) (Fig. 6C). Upregulated pathways along this differentiation route indicate progression to a more ECM catabolic state indicative of reactive/remodeling tissue, including negative regulation of angiogenesis (GO:0016525), collagen fibril organization (GO:0030199), and cell migration (GO:0016477) (Fig 6D). To determine which transcription factors (TFs) may be regulating this differentiation trajectory, we performed the TF binding motif analysis using RcisTarget R package and TRRUST database. The results indicated that *Fosl1, Hmga2*, and *Snai1* are the potential key TFs driving this differentiation process with *Fosl1* and *Snai1* having higher expression levels and percentage of cells in highly reactive remodeling tendon (cluster 4) (Fig. 6E). When mapping these TFs back onto the spatial location at D14 and 28 samples, we observed that *Snai1* was more distributed throughout the tendon and bridging tissues, while *Fosl1* and *Hmga2* were predominately expressed at tendon stub (Fig. 6F & G). Protein-protein network for module 2 was further constructed using String database and proteins having strong interactions with Fosl1, Hmga2, and Snai1 were also identified (Supplemental Figure 8A &B).

The differentiation of non-reactive/ native tendon-like cells (cluster 1) into peripheral fibroblastic cells (cluster 0) was best described by gene module 5 (Fig 7A & B, Supplemental Figure 7). This trajectory showed upregulation of *CD248, Postn, Clec3b*, and *Ptpn18* at the later differentiation stage (i.e., peripheral fibroblastic cells) (Fig. 7C). Interestingly, the most highly enriched pathways in this trajectory include many related to immune response (GO:0002376~immune system response, GO:0045087~innate immune response), as well as inflammatory pathways (GO:0006955~inflammatory response, GO:0030593~neutrophil chemotaxis) (Fig. 7D). TF binding motif analysis shows that the primary TFs regulating the differentiation route are *Spi1, Hif1a*, and *Klf4* (Fig. 7E). The spatial localization analysis indicates that *Spi1, Hif1a*, and *Klf4* had wider distribution in tendon tissue than *Fosl1, Hmga2*, and *Snai1* identified in module 2 at D14 and D28 samples (Fig. 7F & G). Protein-protein network for the module 5 trajectory was also constructed (Supplemental Figure. 9A &B).

**Figure 7.**
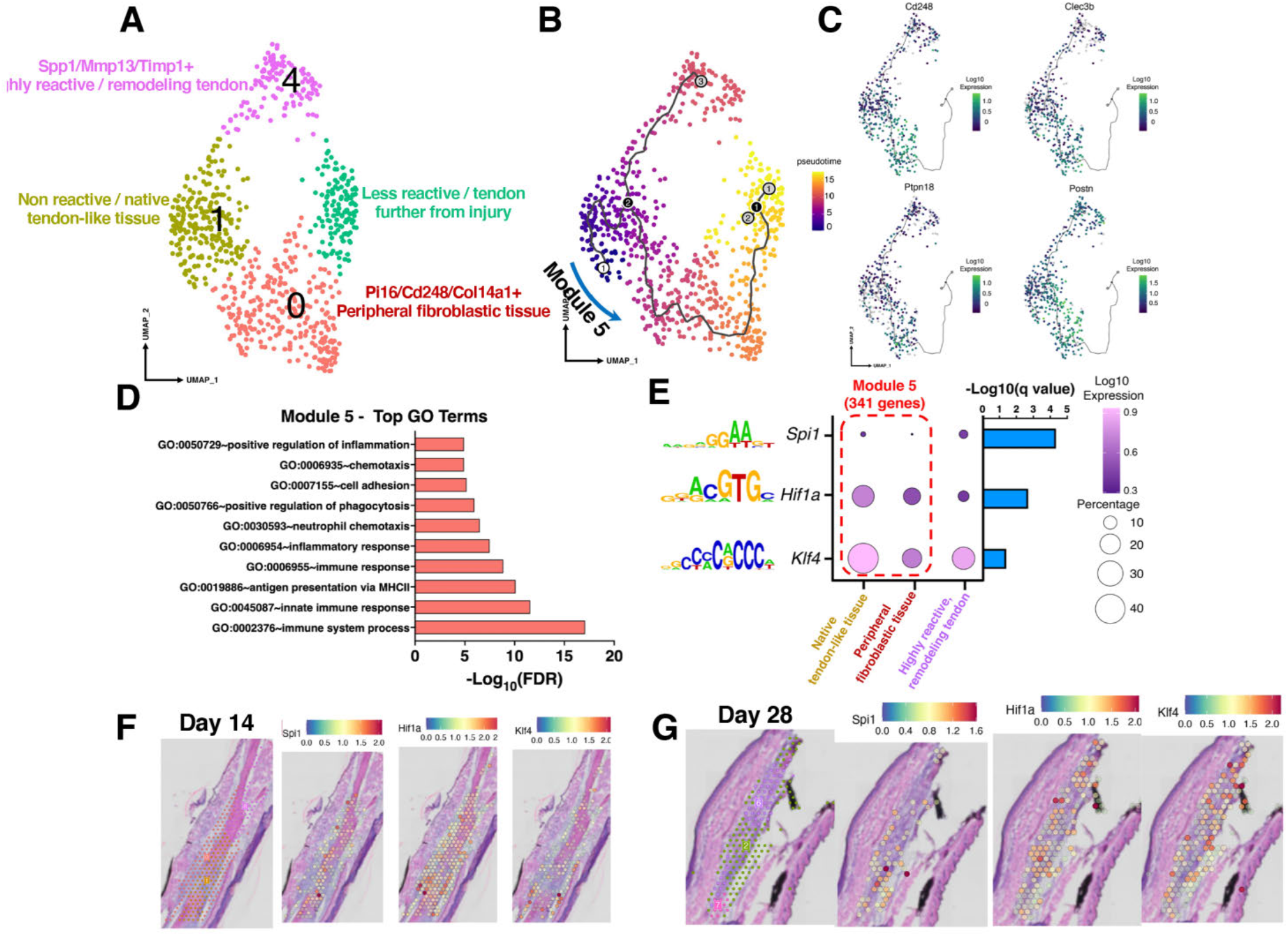
Pseudotime trajectory analysis from basal tendon to scar capsule to less reactive tissue. (A) UMAP of the clusters included in pseudotime analysis (Macrophage cluster was excluded). (B) Pseudotime analysis demonstrating the location of module 5 which was used for subsequent analysis. (C) Identification of genes that define the trajectory in module 5, including *Cd248, Clec3b, Ptpn18*, and *Postn*. (D) Identification of GO terms enriched in module 5 trajectory. (E) TF binding motif analysis predicts transcription factors *Spi1, Hif1a*, and *Klf4* as regulators of the trajectory in module 5. (F & G) Spatial mapping of the subclusters included in pseudotime analysis and the predicted transcription factors regulating this trajectory at D14 (F) and D28 (G).

### Cellular interactome analysis defines crosstalk between macrophages and remodeling tendon tissue

With spatially resolved cell populations and their associated transcriptomic profiles, we can now elucidate cell-cell crosstalk not only within the same cell population but also distinct cell populations between selected areas that are physical proximity to each other. Thus, we next examined the interaction between inflammatory bridging tissue/macrophages (cluster 3) and highly reactive remodeling tendon (cluster 4) to define the signaling pathways that may drive the tendon healing process. Several pathways involved in fibrotic scar formation and inflammation including TGFβ-, EGFR-, IL1R-and TNFR-mediated signaling were identified (Fig. 8). For example, remodeling tendon tissue highly expresses TGFBR3, and that this pathway can be stimulated by inflammatory macrophage secretion of TGFB1 and TGFB3 ligands to drive fibrotic healing (Fig 8). Additionally, CD44, a cell surface glycoprotein involved in cell-cell interactions and adhesion, is primarily expressed by cells in the inflammatory bridging tissue and interacts strongly with HBEGF (heparin-binding EGF-like growth factor) in the reactive tendon tissue (Fig. 8).

**Figure 8.**
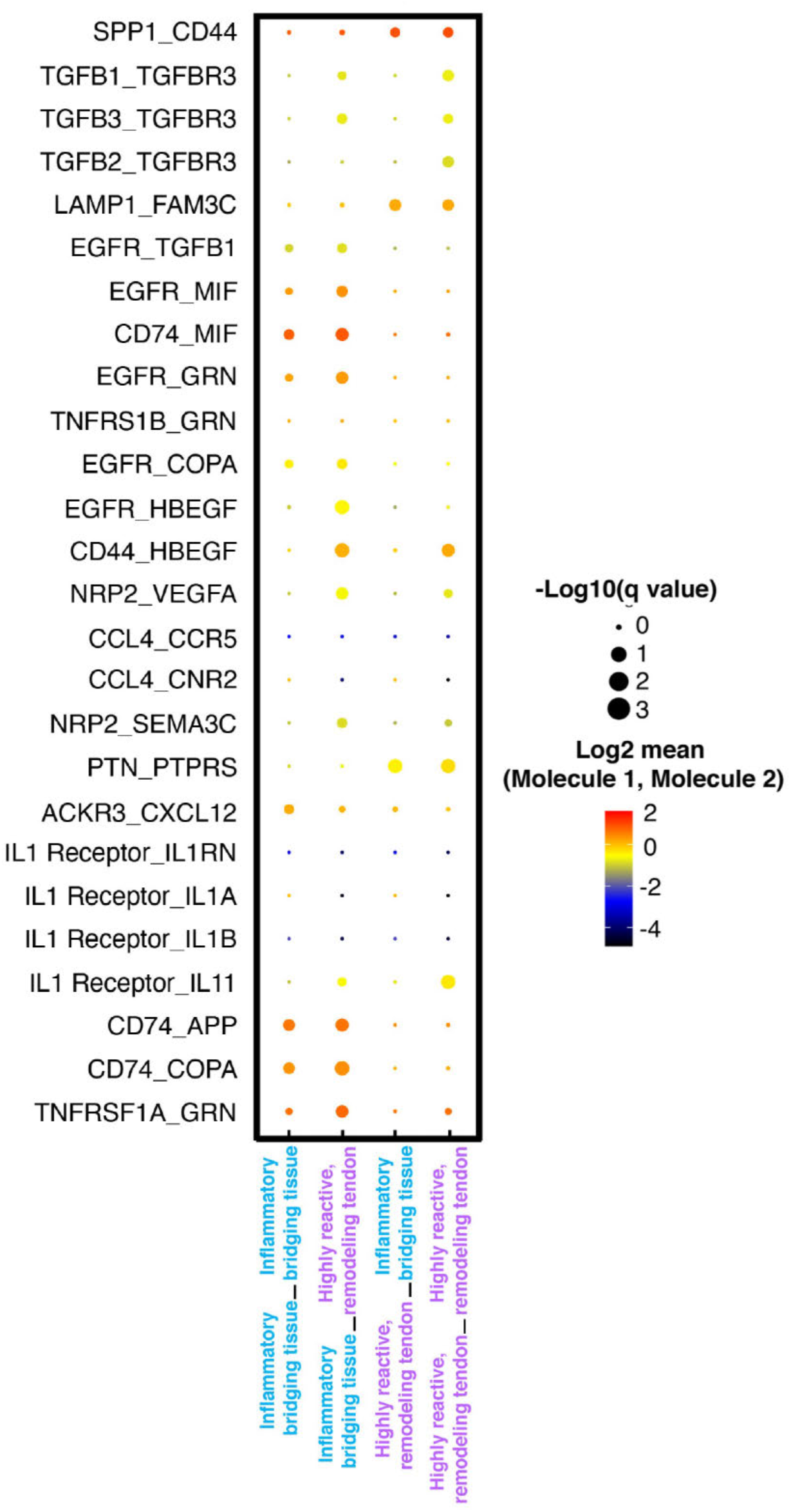
Identification of the cellular interactome at D14. Dot plot of the identified receptor-ligand interactions between the inflammatory bridging tissue (cluster 3) and the highly reactive remodeling tendon (cluster 4) which are known to be in spatial proximity to each other. The first tissue listed in each pair on the x-axis corresponds to the first molecule listed in each pair on the y-axis, while the second tissue on the x-axis, corresponds to the second molecule in each pair on the y-axis. Dot size and color indicate the -Log10 of the qvalue, and the Log2 of the mean, respectively.

## Discussion

Dissecting the molecular programs that underly both physiological and pathological aspects of tendon healing is critical to the development and translation of biological and pharmacological approaches to improve tendon healing. Here we have demonstrated the time- and location-dependent proliferation, activation, myofibroblast differentiation of adult Scx^Lin^ cells (Scx^Ai9^) during tendon healing and defined how additional populations of Scx+ cells are added to the overall Scx^Lin^ pool throughout the healing process. These data have demonstrated a unique spatial response in which Scx^Ai9^ cells initially contribute to an organized cellular bridge at the repair site with an absence of myofibroblast differentiation observed, followed by a progressive differentiation to aSMA+ myofibroblasts, with these cells primarily restricted to the remodeling stubs of the native tendon. Given the importance of the myofibroblast fate to both successful wound healing and fibrotic progression, we sought to define the unique spatio-molecular programs underlying both overall healing, and these distinct responses of Scx^Ai9^ cells, using spatial transcriptomics.

During normal wound healing, ‘fibroblast activation’, in which tissue-resident fibroblast populations break quiescence, proliferate and contribute to the healing is typically initiated via production of damage associated molecular proteins (DAMPs) ^25^, and activation is strongly associated with subsequent myofibroblast differentiation ^26, 27^. While fibroblast activation has been observed in tenocytes ^28, 29^, the cell-lineage of myofibroblast origin is also associated with function ^30, 31^, thus we have focused more specifically on the activation profile and myofibroblast differentiation of adult Scx^Lin^ (Scx^Ai9^) cells. Somewhat surprisingly, we found that activation markers were predominantly associated with Scx^Ai9^ cells in the organized neo-tendon bridge, with these cells undergoing a transient differentiation to aSMA+ myofibroblasts, followed by loss of aSMA staining in these cells by D14. In contrast, minimal expression of activation markers was observed in Scx^Ai9^ in the stubs of native tendon even though these cells progress to a more persistent myofibroblast fate through D28. In addition, combining the adult Scx^Ai9^ trace with the Scx^GFP^ reporter demonstrated clear maintenance of *Scx* expression in myofibroblasts regardless of their adult Scx^Ai9^ origin, suggesting a potentially important role for *Scx* in myofibroblast differentiation. Taken together, these data demonstrate the ability of Scx^Ai9^ cells in the neo-tendon bridge to undergo transient myofibroblast differentiation, although future work will be needed to determine if these cells ‘revert’ to a more basal tenocyte phenotype or are cleared via apoptosis. Moreover, the differences in activation profile between transient and persistent myofibroblast suggest that unique molecular programs likely underlie these fate decisions, and as such represent potentially important intervention opportunities to modulate myofibroblast activity.

With spatial transcriptomic analysis, one of the novel findings was the discrete molecular programs that were observed in the proximal and distal ends of the native tendon at D14. While there was a main cluster that encompassed the bridging neo-tendon and the most reactive tendon stubs immediately adjacent to the repair site, distinct clusters were observed proximal and distal to this cluster. While the reason for this is unclear, it may be due to slight differences in the mechanobiological environment within unique anatomic location of the tendon, with the distal end being relatively close to the bony insertion, while the proximal end is quite distant from the myotendinous junction. In addition to these differences, we also observed a progressive change in the tendon molecular profile as distance from the repair site increased, moving from a highly reactive remodeling tissue (characterized by *Spp1* and *Mmp13*) to a more moderately reactive tissue (*Tnmd, Col1a1*), and then to a more basal tendon state (*Pi16, Clec3b*) far away from the repair site, indicating the relatively focal and localized nature of the tendon response to injury. Interestingly, in the integrated analysis, Cochlin (*Coch*) was identified as a defining gene for the non-reactive/ native tendon-like tissue. *Coch* is an ECM protein that comprises the predominant non-collagen ECM component of the cochlea and vestibule of the inner ear ^32^, and *Coch* mutations lead to sensineuronal healing loss ^33^. While the role of *Coch* in tendon is not clear, recent work from Wunderli *et al*., demonstrates downregulation of Coch in an ex-vivo model of hypervascular/ matrix unloading-mediated tendinopathy ^34^. Moreover, Wang *et al*., demonstrated that *Scx* is required for the formation of tendon in the middle ear ^35^, and Coch expression is observed in these structures ^36^, suggesting a potential role for tenocyte-derived Cochlin in tendon homeostasis.

Using pseudotime analysis we identified two distinct molecular program trajectories (when the inflammatory cluster was excluded): one trajectory moving the native tendon-like cluster toward the highly reactive remodeling tendon (module 2), and the other moving from native tendon-like cluster toward the peripheral fibroblastic shell (module 5). Furthermore, putative TFs potentially governing these two distinct cell fate decisions were also identified in our study. Interestingly, tdTomato (indicative of adult Scx^Ai9^ cells) was a defining feature of the shift of transcriptional profile from basal, native to highly reactive tissue, indicating a central role for these cells in this process, and consistent with their persistent localization within the reactive tissue. In addition, the differentiation trajectory toward highly reactive remodeling tendon cells was associated with GO terms such as wound healing, collagen catabolic processes, and collagen fibril organization, suggesting that adult Scx^Ai9^ cells may modulate healing via these processes. In contrast, tdTomato was not associated with the trajectory in module 5. This trajectory, which was associated with a peripheral fibroblast tissue, skewed more heavily toward inflammatory response including positive regulation of inflammation and immune system processes. While future work will be needed to better understand the potential differential functions of these tissue areas it is notable that the exterior scar tissue shell that encompasses this trajectory is critical in the formation of peritendious adhesions, with this pathology perhaps being mediated by a sustained inflammatory program.

While spatial transcriptomics does not provide the resolution at the single cell levels, one of its major benefits is the ability to interrogate the potential cell-cell crosstalk between clusters that are known to be in physical proximity, thereby allowing identification of potential key interactions underpinning the molecular programs. Our cellular interactome data (of the integrated dataset) focused on the interaction between the inflammatory (macrophage-like) bridging tissue and the highly reactive remodeling tendon at the injury site, as these two clusters are physically adjacent to each other. Several pathways involved in fibrotic scar formation and inflammation including TGFβ-, EGFR-, IL1R- and TNFR-mediated signaling were identified between these two cell populations. Moreover, the CD44-HBEGF interaction from the inflammatory bridging tissue, and the reactive remodeling tendon, respectively, is of particular interest as this interaction is part of a cell surface complex that plays an important role in physiological tissue remodeling and cell survival^37^, suggesting that inflammatory macrophages may mediate tendon remodeling via CD44 interaction with HBEGF in tendon cells.

One of the key limitations to this study is the small sample size for spatial transcriptomic analysis. Nevertheless, no apparent batch effects were observed in two independent D14 samples (biological replicates) in the current study, suggesting that our injury outcomes and sequencing results are highly reproducible. While this study provides significant information regarding spatially-resolved transcriptomic profiles of tendon healing in young, metabolically healthy mice, the effects of aging, or other comorbidities on the molecular program of tendon healing may warrant future investigations.

Defining the distinct molecular programs that predominate during different phases of healing in a spatially dependent way will substantially enhance our understanding of the complex cellular and molecular milieu of tendon healing and will facilitate identification of more targeted therapeutic approaches to improve tendon healing.

## Author contributions

Study conception and design: JEA, KTB, AEL; Acquisition of data: JEA, KTB, SNM; Analysis and interpretation of data: KTB, JEA, CLW, AEL; Drafting of manuscript: JEA, KTB, CLW, AEL; Revision and approval of manuscript: JEA, KTB, SNM, CLW, AEL.

## Acknowledgements

We would like to thank the Histology, Biochemistry and Molecular Imaging (HBMI) Core for technical assistance with the histology. We would also like to thank the UR Genomics Research Core for assistance with Spatial Transcriptomics.

## Supplemental Figures

**Supplemental Figure 1.**
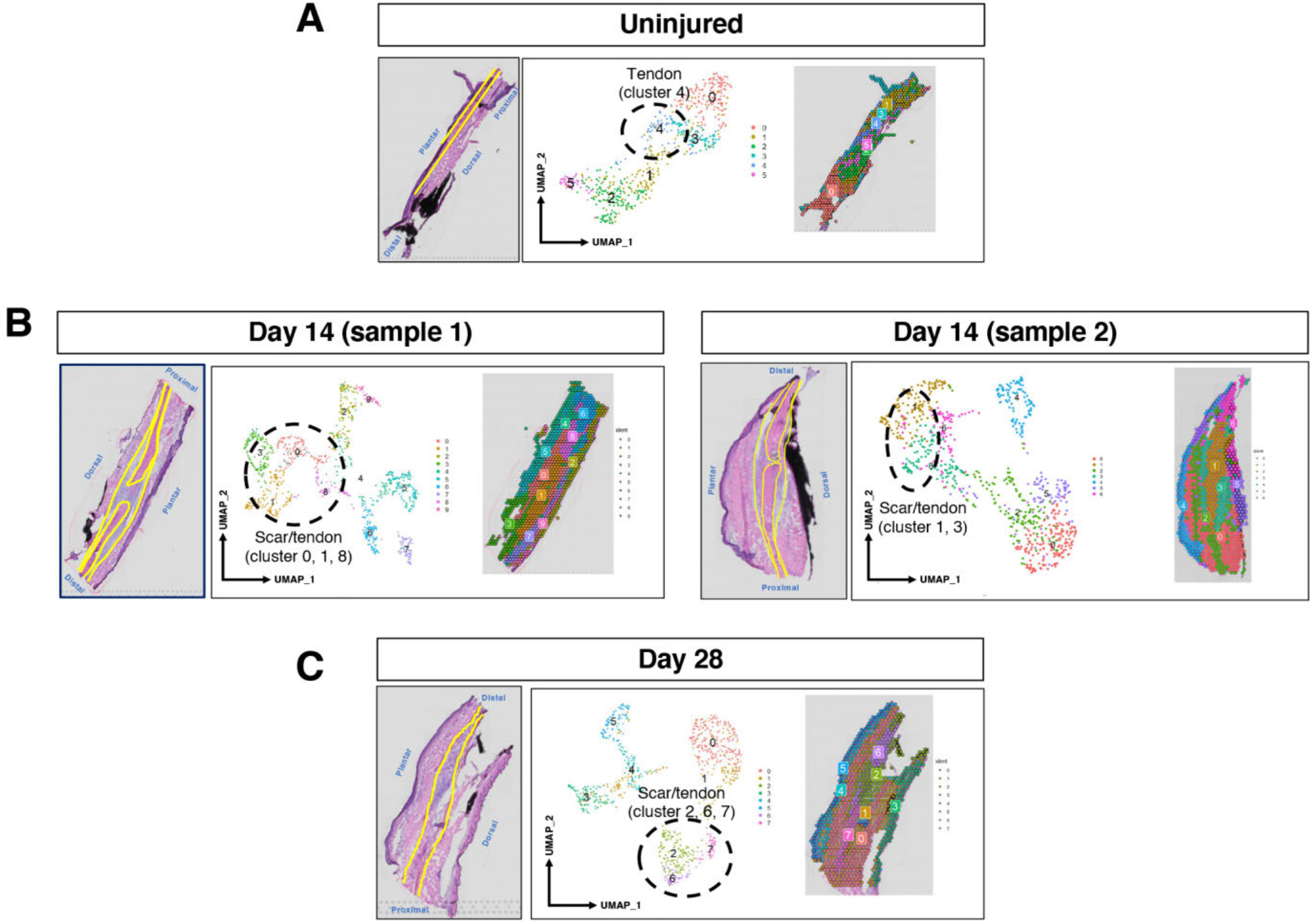
Unsupervised clustering and identification of tendon and scar clusters. (A) Native tendon is outlined in yellow in the histological image of an uninjured tendon, followed a UMAP of the unsupervised clustering of all transcriptomics data captured from that slide, and spatial mapping of the unsupervised clusters. (B & C) Native tendon and scar tissue are outlined in yellow at D14 (B) and D28 (C). All data underwent unsupervised clustering, and the clusters used for subsequent analysis were identified based on their overlap with tendon and scar tissue. Clusters that were used for subsequent analysis are circled on their respective UMAPs.

**Supplemental Figure 2.**
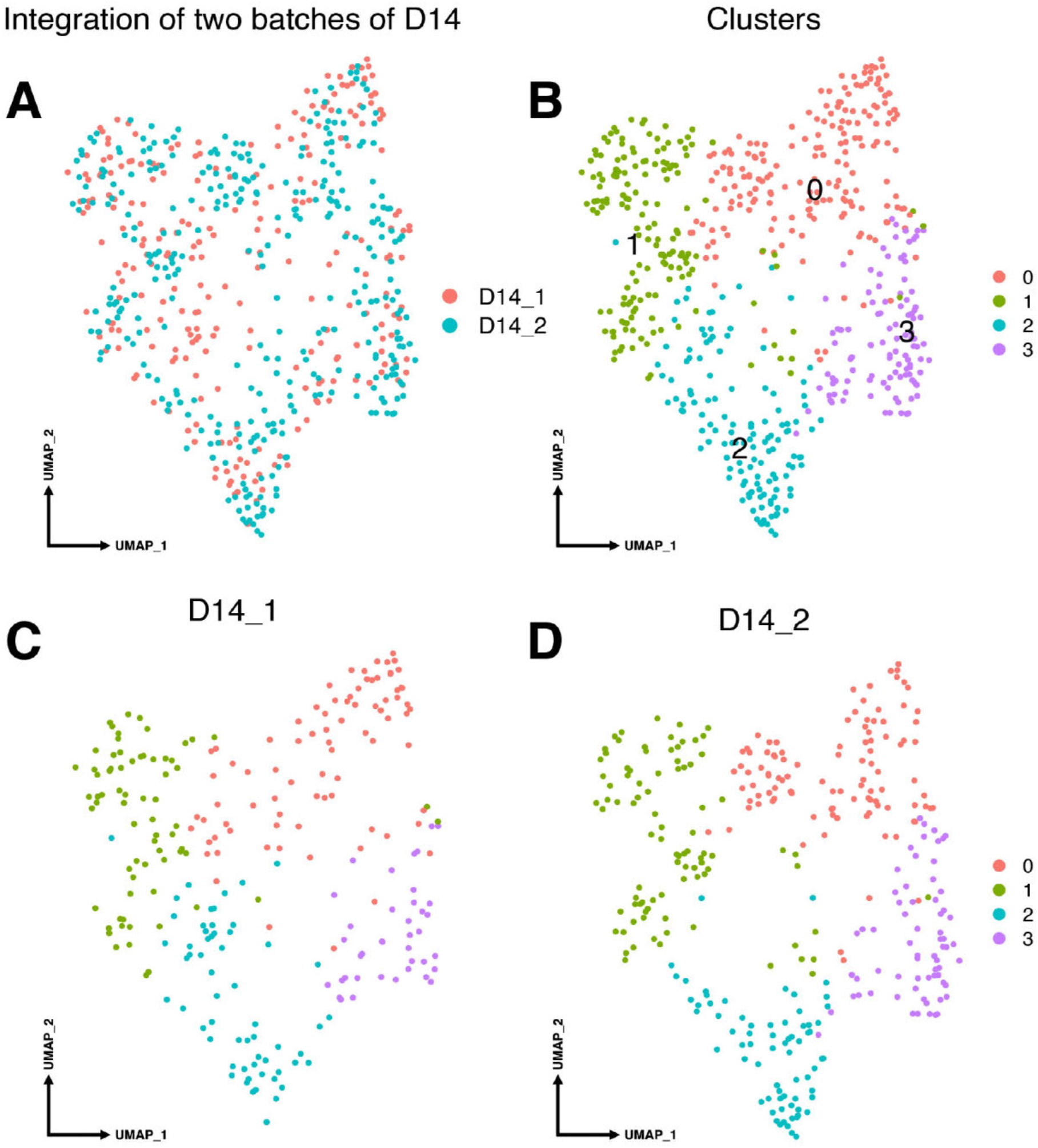
Analysis of batch effects at D14. (A) UMAP of biological replicates, D14_1 is shown in red, and D14_2 in blue. (B) UMAP of unsupervised clustering for the merged dataset demonstrates 4 distinct clusters that are shared in both the D14_1 sample (C), and in D14_2 (D).

**Supplemental Figure 3.**
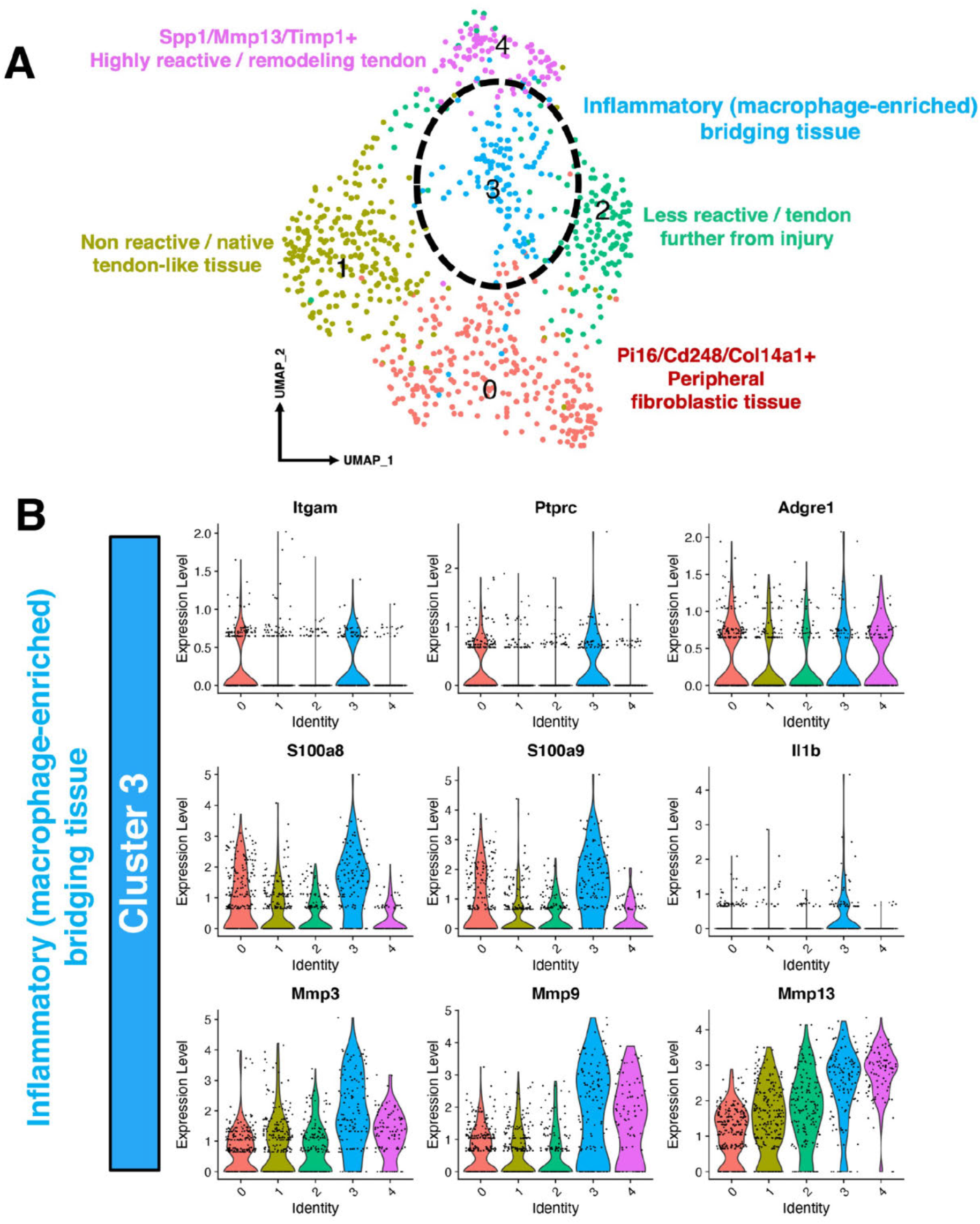
Cluster 3 in the integrated dataset marks an inflammatory bridging tissue population. This cluster is identified in the integrated UMAP in (A), and selected gene expression shown with violin plot in (B). Compared to other clusters, cluster 3 shows high expression levels of *Acp5, Cd11b* (*Itgam*), *Cd45* (*Ptprc*), F4/80 (*Adgre1*) the alarmins *S100a8, S100a9*, and *IL1b*, as well as matrix remodeling genes *Mmp9* and *Mmp13*.

**Supplemental Figure 4.**
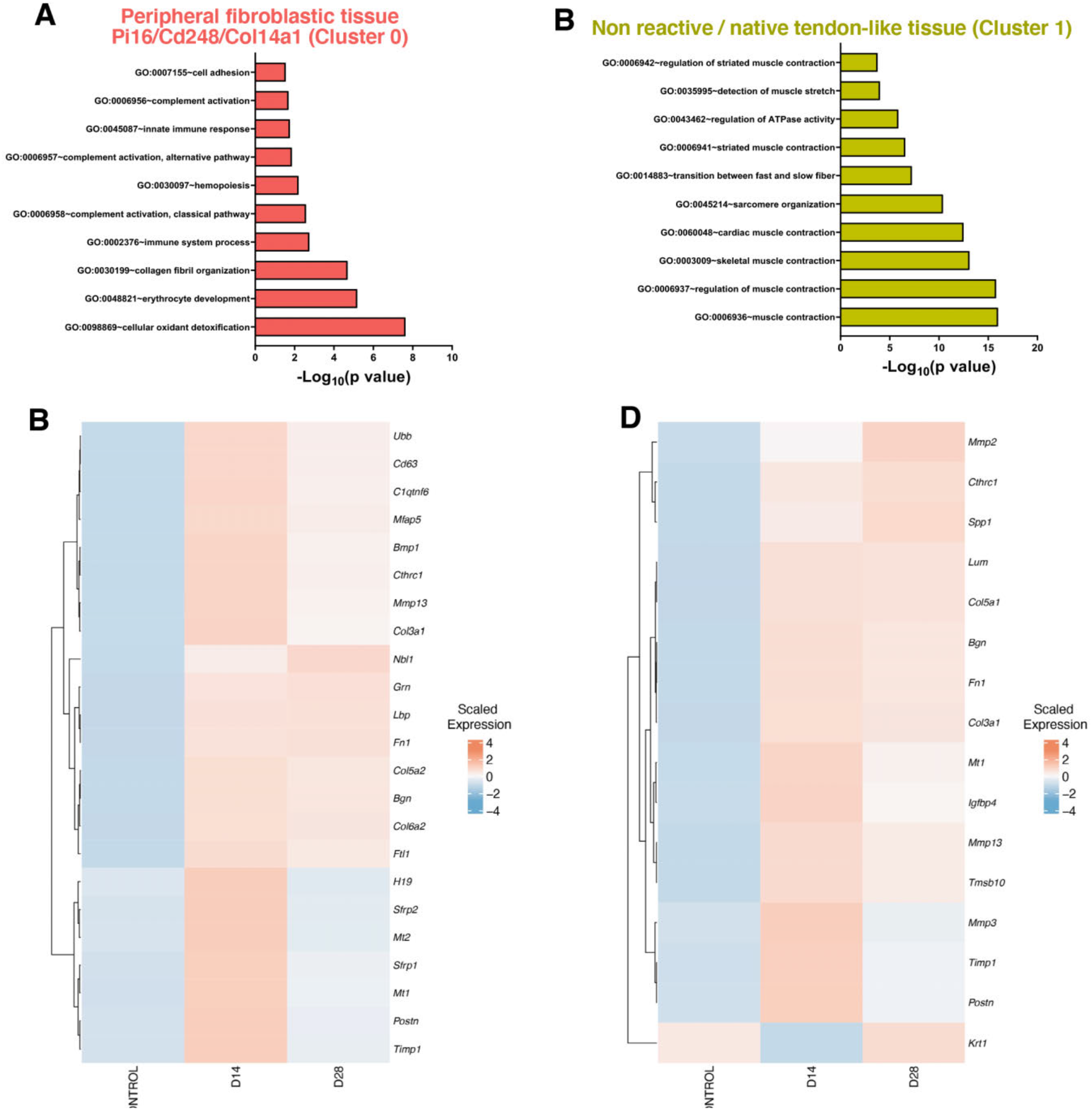
GO term analysis and heatmap showing DEGs over time for clusters 0 and 1. Significantly upregulated pathways in the peripheral fibroblastic tissue Pi16/Cd248/Col14a1 (Cluster 0) are shown in (A), along with a heatmap of selected genes in this cluster over time (control = uninjured) in (B). Pathways in the non reactive / native tendon-like tissue (Cluster 1) are shown in (C), with corresponding heatmap in (D).

**Supplemental Figure 5.**
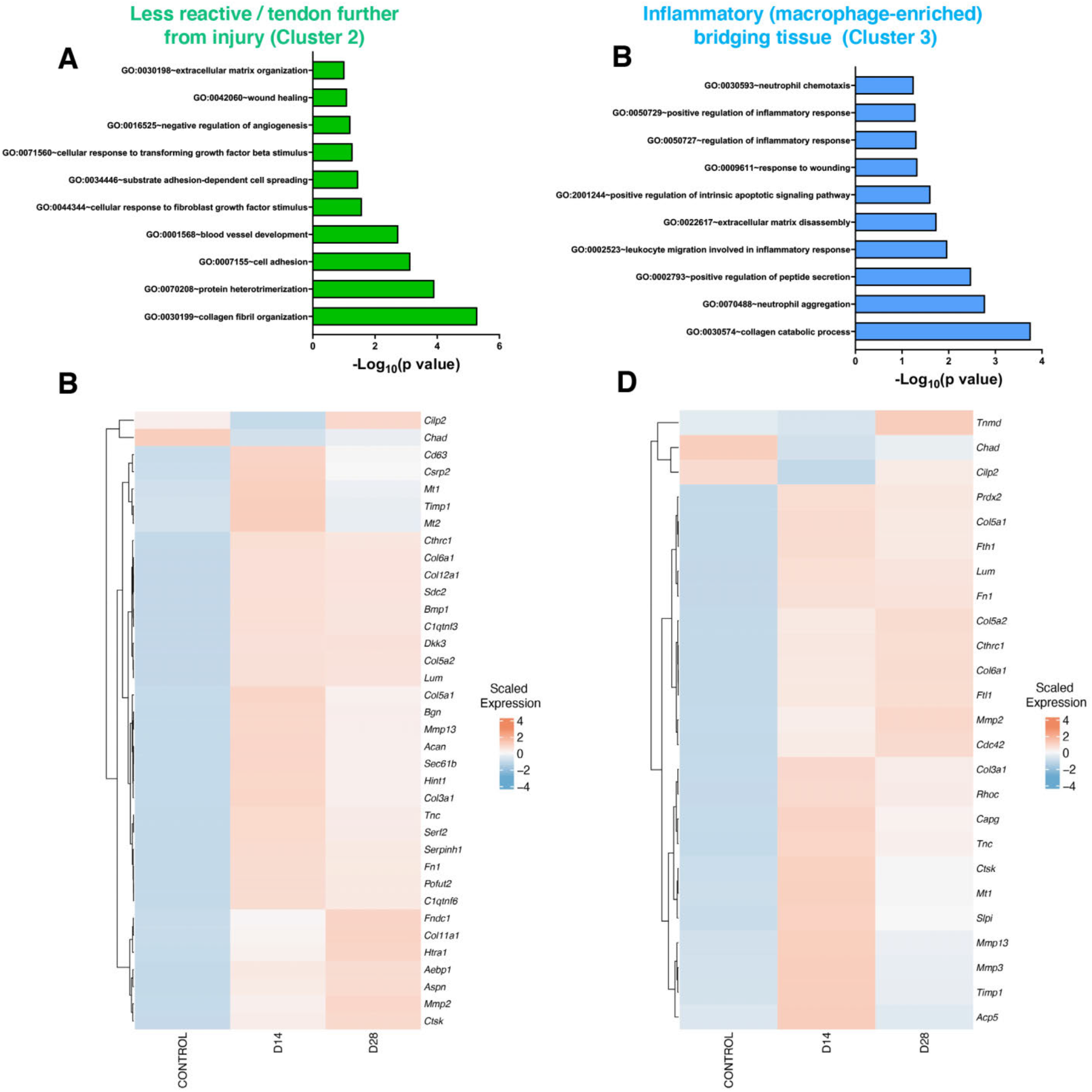
GO term analysis and heatmap showing DEGs over time for clusters 2 and 3. The less reactive / tendon further from injury (Cluster 2) demonstrated significant upregulation of pathways shown in (A), and heatmap of this cluster over time (control = uninjured) is shown in (B). Cluster 3, annotated as the inflammatory (macrophage-enriched) bridging tissue, exhibited upregulation of the pathways shown in (C), with a corresponding heaatmap in (D).

**Supplemental Figure 6.**
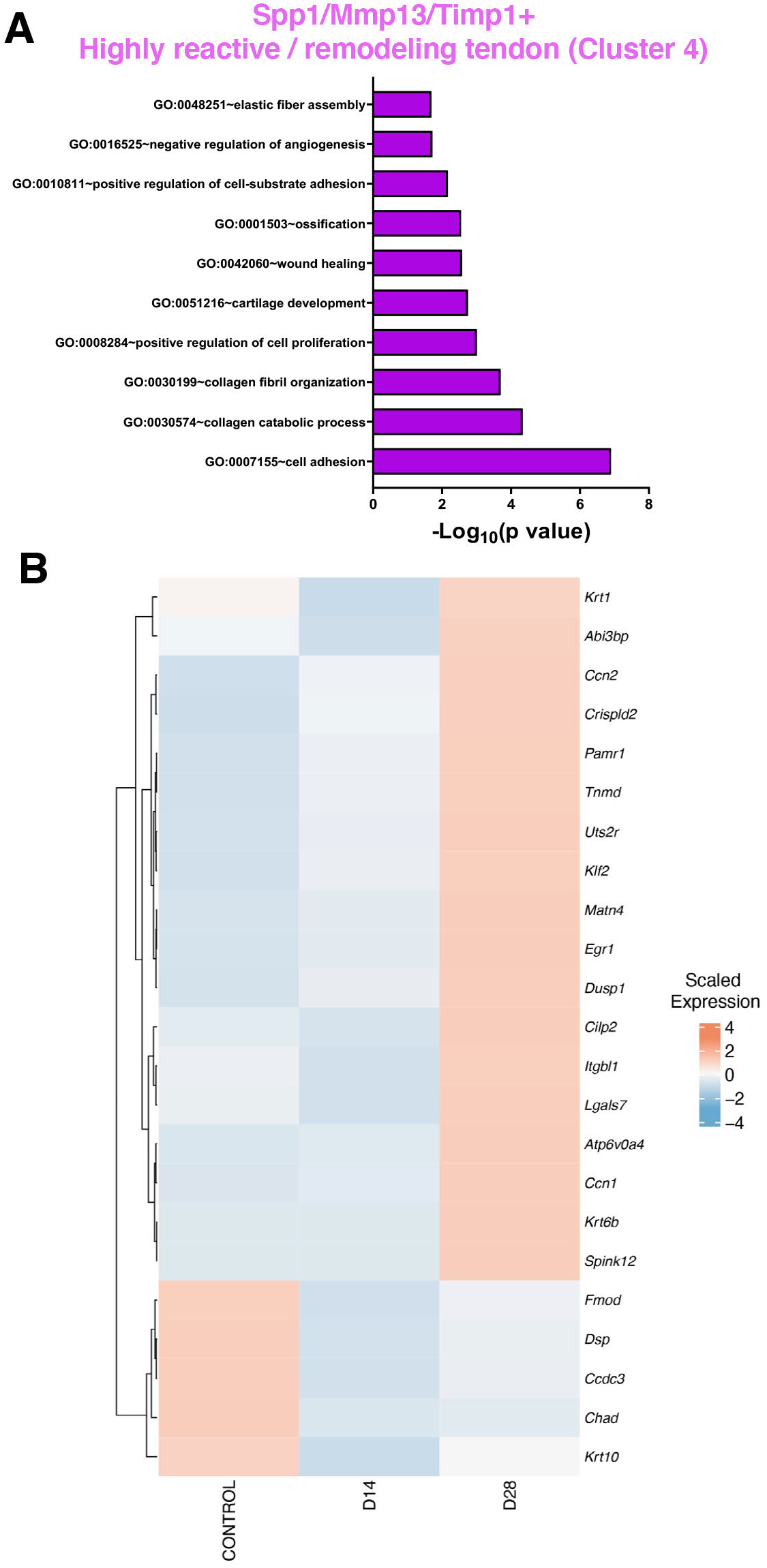
GO term analysis and heatmap showing DEGs over time for cluster 4. Significantly upregulated pathways in the Spp1/Mmp13/Timp1+ highly reactive / remodeling tendon (Cluster 4), are shown in (A), and heatmap of selected genes for this cluster over time in (B).

**Supplemental Figure 7.**
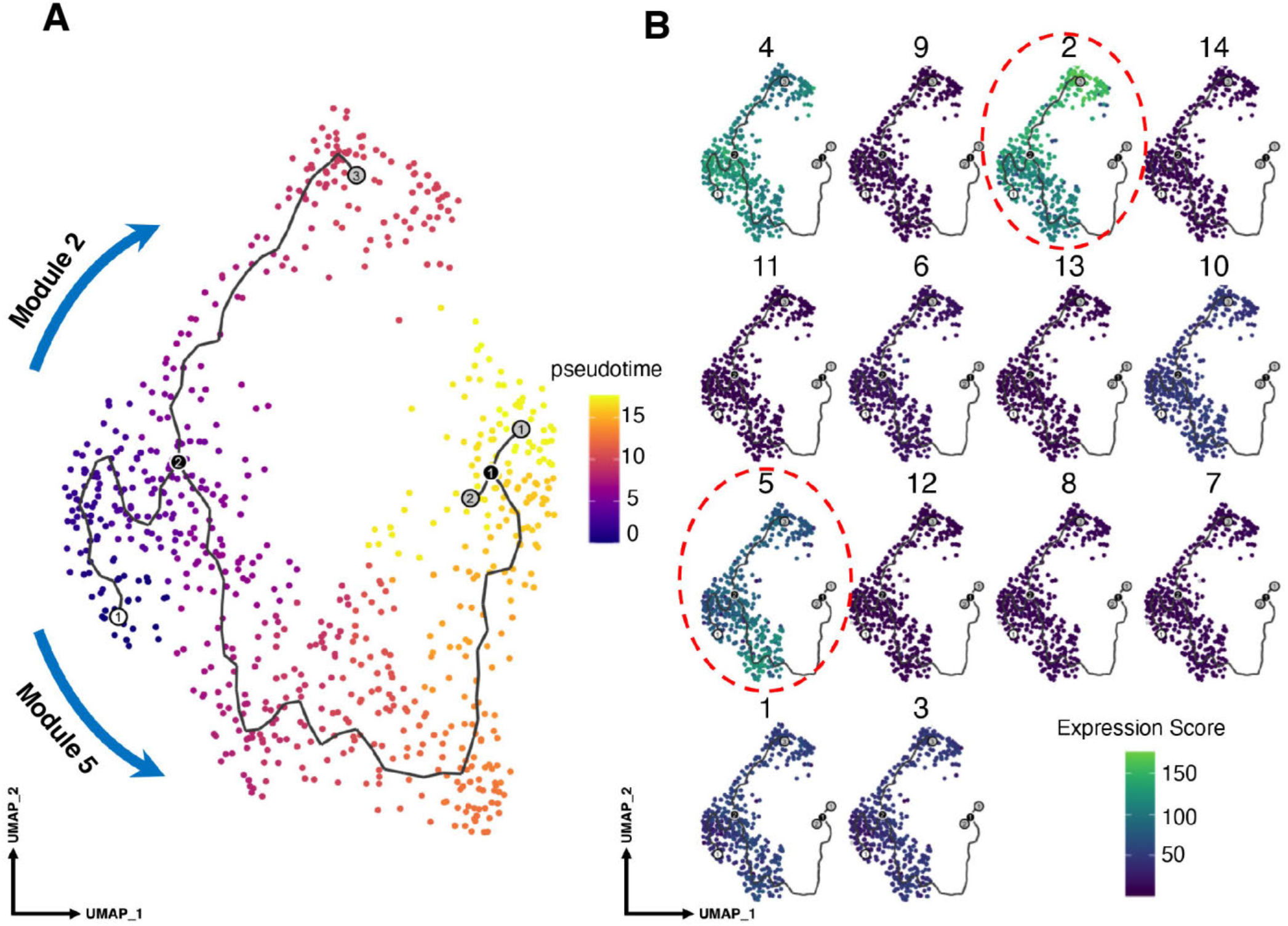
Gene module analysis of differentiation trajectories. The Mmp3/Mmp9 enriched bridging tissue cluster was first removed from the integrated spatial-RNA-seq dataset (see *Methods*), and potential differentiation trajectories in our integrated spatial-seq dataset identified in (A), using the non-reactive/ native tendon-like cell cluster (1) as the starting point. Gene modules were generated to describe each trajectory using hierarchical cluster analysis in (B), and gene module 2 best defined cell fate decisions of non-reactive/ native tendon-like cells into highly reactive remodeling tendon cells, while genes in module 5 regulated the differentiation of non-reactive/native tendon-like cells into peripheral fibroblastic cells.

**Supplemental Figure 8.**
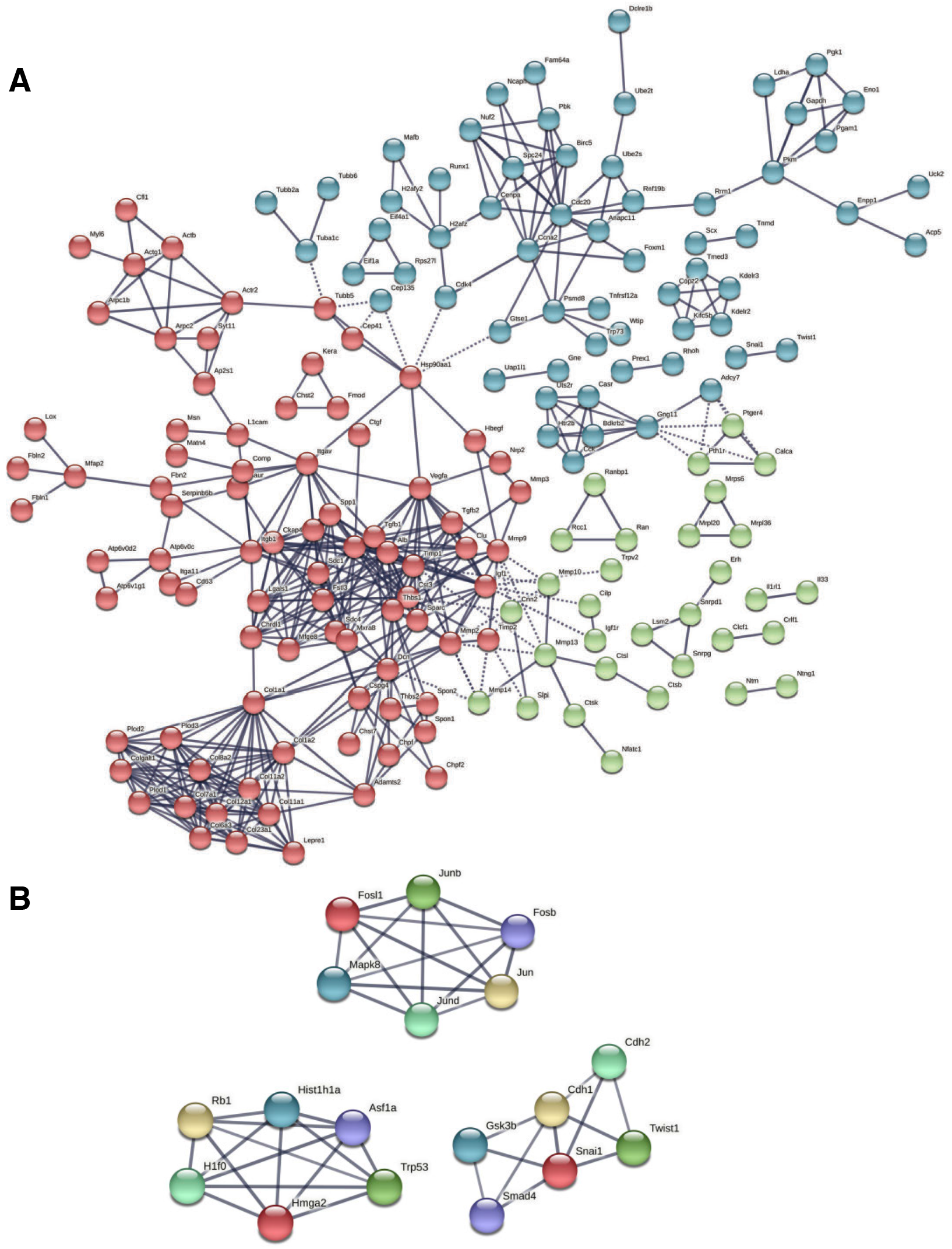
Module 2 protein-protein network analysis. Genes regulating cell fate decisions of non-reactive/ native tendon-like cells to highly reactive remodeling tendon cells were matched to a known database of protein-protein interactions (see *Methods*), creating the network shown in (A). This constructed network was then further clustered with k-means clustering (B) to include key TF described in Figure 7.

**Supplemental Figure 9.**
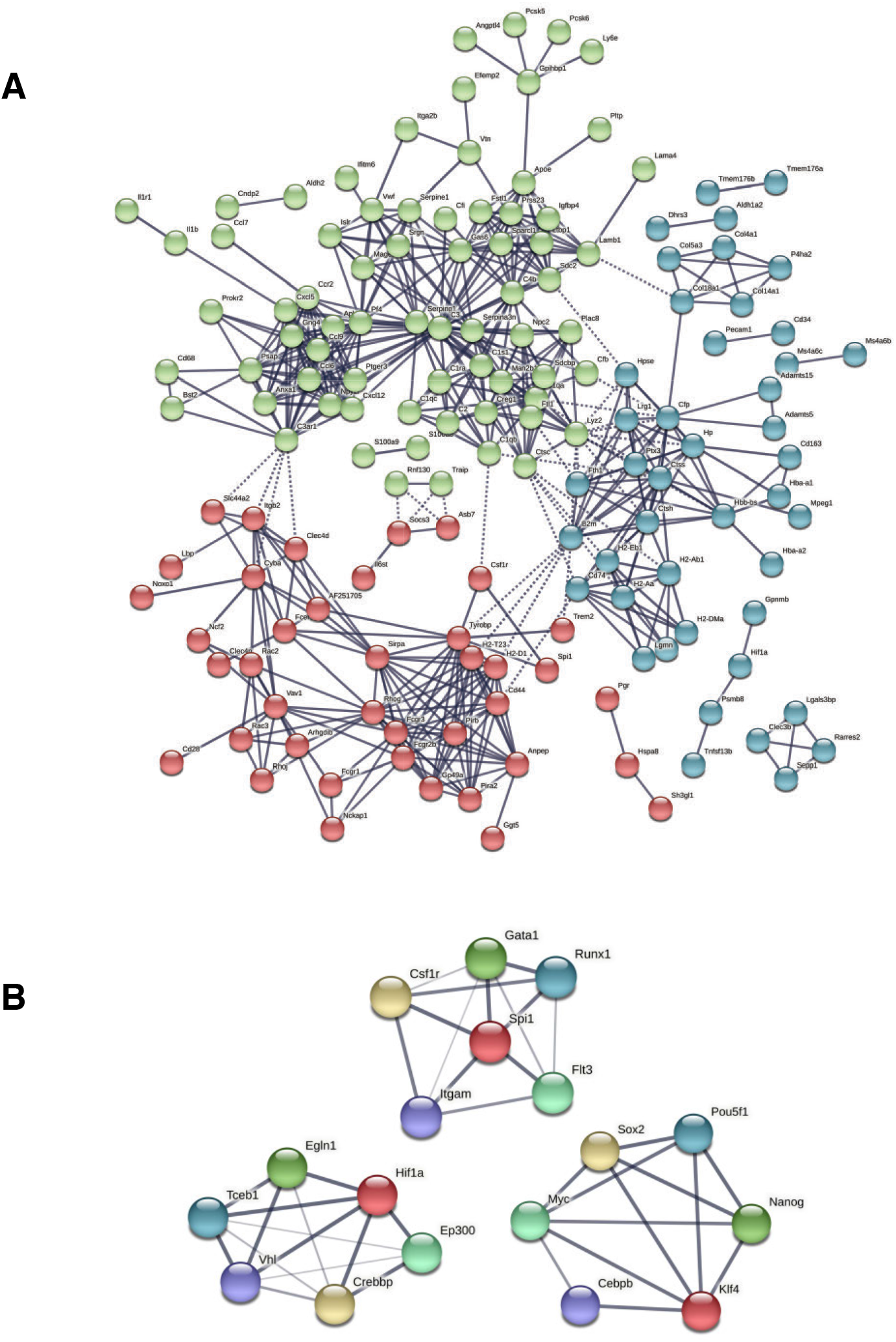
Module 5 protein-protein network analysis. Protein-protein interactions regulating the differentiation trajectory of non-reactive *I* native tendon-like cells to the peripheral fibroblastic tissue (Pi16/Cd248/Col14a1+) are shown in (A), along with k-means clustering (B) describing the network surrounding main TF described in Figure 8.

## References

1. Murchison, N.D., B.A. Price, D.A. Conner, D.R. Keene, E.N. Olson, C.J. Tabin, and R. Schweitzer, Regulation of tendon differentiation by scleraxis distinguishes force-transmitting tendons from muscle-anchoring tendons. Development, 2007.

2. Yoshimoto, Y., A. Takimoto, H. Watanabe, Y. Hiraki, G. Kondoh, and C. Shukunami, Scleraxis is required for maturation of tissue domains for proper integration of the musculoskeletal system. Sci Rep, 2017. 7: p. 45010. 10.1038/srep45010.

3. Dyment, N.A., Y. Hagiwara, B.G. Matthews, Y. Li, I. Kalajzic, and D.W. Rowe, Lineage tracing of resident tendon progenitor cells during growth and natural healing. PLoS One, 2014. 9(4): p. e96113. 10.1371/journal.pone.0096113.

4. Howell, K., C. Chien, R. Bell, D. Laudier, S.F. Tufa, D.R. Keene, N. Andarawis-Puri, and A.H. Huang, Novel Model of Tendon Regeneration Reveals Distinct Cell Mechanisms Underlying Regenerative and Fibrotic Tendon Healing. Sci Rep, 2017. 7: p. 45238. 10.1038/srep45238.

5. Best, K.T., A. Korcari, K.E. Mora, A.E. Nichols, S.N. Muscat, E. Knapp, M.R. Buckley, and A.E. Loiselle, Scleraxis-lineage cell depletion improves tendon healing and disrupts adult tendon homeostasis. Elife, 2021. 10. 10.7554/eLife.62203.

6. Best, K.T. and A.E. Loiselle, Scleraxis lineage cells contribute to organized bridging tissue during tendon healing and identify a subpopulation of resident tendon cells. FASEB J, 2019. 33(7): p. 8578–8587. 10.1096/fj.201900130RR.

7. Pakshir, P. and B. Hinz, The big five in fibrosis: Macrophages, myofibroblasts, matrix, mechanics, and miscommunication. Matrix Biol, 2018. 68-69: p. 81–93. 10.1016/j.matbio.2018.01.019.

8. Loiselle, A.E., G.A. Bragdon, J.A. Jacobson, S. Hasslund, Z.E. Cortes, E.M. Schwarz, D.J. Mitten, H.A. Awad, and R.J. O’Keefe, Remodeling of murine intrasynovial tendon adhesions following injury: MMP and neotendon gene expression. J Orthop Res, 2009. 27(6): p. 833–40. 10.1002/jor.20769.

9. Ackerman, J.E. and A.E. Loiselle, Murine Flexor Tendon Injury and Repair Surgery. J Vis Exp, 2016(115). 10.3791/54433.

10. Hao, Y., S. Hao, E. Andersen-Nissen, W.M. Mauck, S. Zheng, A. Butler, M.J. Lee, A.J. Wilk, C. Darby, and M. Zagar, Integrated analysis of multimodal single-cell data. bioRxiv, 2020.

11. Becht, E., L. McInnes, J. Healy, C.A. Dutertre, I.W.H. Kwok, L.G. Ng, F. Ginhoux, and E.W. Newell, Dimensionality reduction for visualizing single-cell data using UMAP. Nat Biotechnol, 2018. 10.1038/nbt.4314.

12. Sherman, B.T., Q. Tan, J.R. Collins, W.G. Alvord, J. Roayaei, R. Stephens, M.W. Baseler, H.C. Lane, and R.A. Lempicki, The DAVID Gene Functional Classification Tool: a novel biological module-centric algorithm to functionally analyze large gene lists. Genome biology, 2007. 8(9): p. 1–16.

13. Gu, Z., R. Eils, and M. Schlesner, Complex heatmaps reveal patterns and correlations in multidimensional genomic data. Bioinformatics, 2016. 32(18): p. 2847–2849.

14. Trapnell, C., D. Cacchiarelli, J. Grimsby, P. Pokharel, S. Li, M. Morse, N.J. Lennon, K.J. Livak, T.S. Mikkelsen, and J.L. Rinn, The dynamics and regulators of cell fate decisions are revealed by pseudotemporal ordering of single cells. Nature biotechnology, 2014. 32(4): p. 381.

15. Qiu, X., Q. Mao, Y. Tang, L. Wang, R. Chawla, H.A. Pliner, and C. Trapnell, Reversed graph embedding resolves complex single-cell trajectories. Nature methods, 2017. 14(10): p. 979.

16. Aibar, S., C.B. González-Blas, T. Moerman, H. Imrichova, G. Hulselmans, F. Rambow, J.-C. Marine, P. Geurts, J. Aerts, and J. van den Oord, SCENIC: single-cell regulatory network inference and clustering. Nature methods, 2017. 14(11): p. 1083–1086.

17. Szklarczyk, D., A.L. Gable, D. Lyon, A. Junge, S. Wyder, J. Huerta-Cepas, M. Simonovic, N.T. Doncheva, J.H. Morris, and P. Bork, STRING v11: protein–protein association networks with increased coverage, supporting functional discovery in genome-wide experimental datasets. Nucleic acids research, 2019. 47(D1): p. D607–D613.

18. Efremova, M., M. Vento-Tormo, S.A. Teichmann, and R. Vento-Tormo, CellPhoneDB: inferring cell–cell communication from combined expression of multi-subunit ligand–receptor complexes. Nature protocols, 2020. 15(4): p. 1484–1506.

19. Howe, K.L., P. Achuthan, J. Allen, J. Allen, J. Alvarez-Jarreta, M.R. Amode, I.M. Armean, A.G. Azov, R. Bennett, and J. Bhai, Ensembl 2021. Nucleic Acids Research, 2021. 49(D1): p. D884–D891.

20. Grinstein, M., H.L. Dingwall, L.D. O’Connor, K. Zou, T.D. Capellini, and J.L. Galloway, A distinct transition from cell growth to physiological homeostasis in the tendon. Elife, 2019. 8. 10.7554/eLife.48689.

21. Sakabe, T., K. Sakai, T. Maeda, A. Sunaga, N. Furuta, R. Schweitzer, T. Sasaki, and T. Sakai, Transcription factor scleraxis vitally contributes to progenitor lineage direction in wound healing of adult tendon in mice. J Biol Chem, 2018. 293(16): p. 5766–5780. 10.1074/jbc.RA118.001987.

22. Dyment, N.A., C.F. Liu, N. Kazemi, L.E. Aschbacher-Smith, K. Kenter, A.P. Breidenbach, J.T. Shearn, C. Wylie, D.W. Rowe, and D.L. Butler, The paratenon contributes to scleraxis-expressing cells during patellar tendon healing. PLoS One, 2013. 8(3): p. e59944. 10.1371/journal.pone.0059944.

23. Best, K.T., A.E.C. Nichols, E. Knapp, W.C. Hammert, C. Ketonis, J.H. Jonason, H.A. Awad, and A.E. Loiselle, NF-kappaB activation persists into the remodeling phase of tendon healing and promotes myofibroblast survival. Sci Signal, 2020. 13(658). 10.1126/scisignal.abb7209.

24. Harvey, T., S. Flamenco, and C.M. Fan, A Tppp3(+)Pdgfra(+) tendon stem cell population contributes to regeneration and reveals a shared role for PDGF signalling in regeneration and fibrosis. Nat Cell Biol, 2019. 21(12): p. 1490–1503. 10.1038/s41556-019-0417-z.

25. Turner, N.A., Inflammatory and fibrotic responses of cardiac fibroblasts to myocardial damage associated molecular patterns (DAMPs). J Mol Cell Cardiol, 2016. 94: p. 189–200. 10.1016/j.yjmcc.2015.11.002.

26. Hinz, B., S.H. Phan, V.J. Thannickal, A. Galli, M.L. Bochaton-Piallat, and G. Gabbiani, The myofibroblast: one function, multiple origins. Am J Pathol, 2007. 170(6): p. 1807–16. 10.2353/ajpath.2007.070112.

27. Hinz, B., Formation and function of the myofibroblast during tissue repair. J Invest Dermatol, 2007. 127(3): p. 526–37. 10.1038/sj.jid.5700613.

28. Dakin, S.G., C.D. Buckley, M.H. Al-Mossawi, R. Hedley, F.O. Martinez, K. Wheway, B. Watkins, and A.J. Carr, Persistent stromal fibroblast activation is present in chronic tendinopathy. Arthritis Res Ther, 2017. 19(1): p. 16. 10.1186/s13075-016-1218-4.

29. Akbar, M., M. McLean, E. Garcia-Melchor, L.A. Crowe, P. McMillan, U.G. Fazzi, D. Martin, A. Arthur, J.H. Reilly, I.B. McInnes, and N.L. Millar, Fibroblast activation and inflammation in frozen shoulder. PLoS One, 2019. 14(4): p. e0215301. 10.1371/journal.pone.0215301.

30. Shook, B.A., R.R. Wasko, G.C. Rivera-Gonzalez, E. Salazar-Gatzimas, F. Lopez-Giraldez, B.C. Dash, A.R. Munoz-Rojas, K.D. Aultman, R.K. Zwick, V. Lei, J.L. Arbiser, K. Miller-Jensen, D.A. Clark, H.C. Hsia, and V. Horsley, Myofibroblast proliferation and heterogeneity are supported by macrophages during skin repair. Science, 2018. 362(6417). 10.1126/science.aar2971.

31. Lynch, M.D. and F.M. Watt, Fibroblast heterogeneity: implications for human disease. J Clin Invest, 2018. 128(1): p. 26–35. 10.1172/JCI93555.

32. Robertson, N.G., A.B. Skvorak, Y. Yin, S. Weremowicz, K.R. Johnson, K.A. Kovatch, J.F. Battey, F.R. Bieber, and C.C. Morton, Mapping and characterization of a novel cochlear gene in human and in mouse: a positional candidate gene for a deafness disorder, DFNA9. Genomics, 1997. 46(3): p. 345–54. 10.1006/geno.1997.5067.

33. Robertson, N.G., J.T. O’Malley, C.A. Ong, A.B. Giersch, J. Shen, K.M. Stankovic, and C.C. Morton, Cochlin in normal middle ear and abnormal middle ear deposits in DFNA9 and Coch (G88E/G88E) mice. J Assoc Res Otolaryngol, 2014. 15(6): p. 961–74. 10.1007/s10162-014-0481-9.

34. Wunderli, S.L., U. Blache, A. Beretta Piccoli, B. Niederöst, C.N. Holenstein, F.S. Passini, U. Silván, L. Bundgaard, U. Auf dem Keller, and J.G. Snedeker, Tendon response to matrix unloading is determined by the patho-physiological niche. Matrix Biol, 2020. 89: p. 11–26. 10.1016/j.matbio.2019.12.003.

35. Wang, L., C.S. Bresee, H. Jiang, W. He, T. Ren, R. Schweitzer, and J.V. Brigande, Scleraxis is required for differentiation of the stapedius and tensor tympani tendons of the middle ear. J Assoc Res Otolaryngol, 2011. 12(4): p. 407–21. 10.1007/s10162-011-0264-5.

36. Robertson, N.G., S.M. Jones, T.A. Sivakumaran, A.B. Giersch, S.A. Jurado, L.M. Call, C.E. Miller, S.F. Maison, M.C. Liberman, and C.C. Morton, A targeted Coch missense mutation: a knock-in mouse model for DFNA9 late-onset hearing loss and vestibular dysfunction. Hum Mol Genet, 2008. 17(21): p. 3426–34. 10.1093/hmg/ddn236.

37. Yu, W.H., J.F. Woessner, Jr., J.D. McNeish, and I. Stamenkovic, CD44 anchors the assembly of matrilysin/MMP-7 with heparin-binding epidermal growth factor precursor and ErbB4 and regulates female reproductive organ remodeling. Genes Dev, 2002. 16(3): p. 307–23. 10.1101/gad.925702.

